# Genome-wide DNA supercoiling arises from transcription and SMC activity and mediates transcriptional negative feedback

**DOI:** 10.64898/2026.03.26.714539

**Authors:** Linying Zhu, Qian Yao, Charan Vemuri, Chongyi Chen

**Author notes:** These authors contributed equally (Linying Zhu, Qian Yao).

## Abstract

Genome-wide DNA supercoiling is closely linked to chromatin organization and gene expression, yet the mechanisms establishing genome-scale supercoiling in living cells and its functional consequences remain unclear. Here, we show that genome-wide supercoiling arises from transcription-driven asymmetric topological relaxation together with contributions from SMC complexes in human cells. During RNA polymerase elongation, human topoisomerases preferentially relax positive over negative supercoils, leading to the accumulation of negative supercoiling around genes. This imbalance enriches supercoiling at transcriptionally active regions including TAD boundaries, and promotes the emergence of large-scale topology. In parallel, SMC complexes independently shape genome-wide supercoiling, with cohesin contributing to interphase topology and condensin establishing an overall positively supercoiled mitotic genome. Functionally, the accumulation of transcription-driven negative supercoiling represses local transcription, revealing a supercoiling-mediated negative feedback mechanism. Together, these findings define the mechanistic basis of genome-scale supercoiling in human cells and establish DNA topology as an integral regulatory layer of transcription.

## Introduction

DNA supercoiling is an intrinsic topological property of chromosomes generated by transcription, replication, and higher-order chromosome organization [1]. DNA topoisomerases, the principal regulators of DNA topology, dynamically resolve supercoiling during these processes[1]. While local supercoiling at individual genes has been extensively studied [2–4], recent quantitative genome-wide measurements have revealed the presence of large-scale DNA supercoiling across the human genome [5]. These findings suggest that DNA topology is present and organized at the genome scale, yet the mechanisms that establish and maintain genome-wide DNA supercoiling in living cells remain unresolved, and its potential impacts on genome regulation and function remain unexplored.

In particular, how chromatin processes and topoisomerase activity collectively establish genome-scale supercoiling is unknown. On one hand, although transcription-generated negative and positive twin-supercoiled domains are well established [6], and negative supercoiling [7–9] or both negative and positive twin-supercoiled domains have been directly observed around genes in living cells, including bacteria [10], yeast [11], and human cells [5], whether and how such local supercoiling accumulation contribute to large-scale genome-wide supercoiling is unclear. On the other hand, while SMC complexes have been proposed to influence DNA topology in vitro [12, 13], direct evidence for their contribution to genome-wide supercoiling in living cells is lacking. Most importantly, whether genome-wide supercoiling is merely a passive byproduct of local transcription or an active regulatory layer for genome function remains unknown.

Here, we investigate the mechanisms underlying the formation and maintenance of genome-wide supercoiling. Using primarily newly generated sequencing data with biological replicates (Table 1), complemented by re-analysis of selected datasets from our previous study [5], we demonstrate that DNA supercoiling across the human genome is established by transcription-driven asymmetric topological relaxation, together with parallel contributions from SMC complexes. We further show that negative supercoiling around genes imposes an inhibitory feedback that constrains transcriptional output, establishing genome-wide DNA topology as an active component of genome regulation.

**Table 1.** Sequencing datasets used in this study. “New” indicates data newly generated for this study, whereas “published” indicates previously published sequencing data that were re-analyzed in this study.

| Sample name | Category | New? | Sample name | Category | New? | Sample name | Category | New? |
| --- | --- | --- | --- | --- | --- | --- | --- | --- |
| HCT116-ATMP-rep1 | ATMP-seq | new | FLV+BRM-rep1 | ATMP-seq | new | TT-HCT116-batch1-rep1 | TT-seq | new |
| HCT116-ATMP-rep2 | ATMP-seq | new | FLV+BRM-rep2 | ATMP-seq | new | TT-HCT116-batch1-rep2 | TT-seq | new |
| HCT116-Ctrl-rep1 | ATMP-seq | new | FLV+TSA-rep1 | ATMP-seq | new | TT-HCT116-batch2-rep3 | TT-seq | new |
| HCT116-Ctrl-rep2 | ATMP-seq | new | FLV+TSA-rep2 | ATMP-seq | new | TT-HCT116-batch3-rep4 | TT-seq | new |
| TPL-rep1 | ATMP-seq | new | FLV+RAD21-IAA-rep1 | ATMP-seq | new | TT-FLV-batch1-rep1 | TT-seq | new |
| TPL-rep2 | ATMP-seq | new | FLV+RAD21-IAA-rep2 | ATMP-seq | new | TT-FLV-batch1-rep2 | TT-seq | new |
| FLV-rep1 | ATMP-seq | new | FLV+ICRF+CPT | ATMP-seq | new | TT-TOP1-IAA+ICRF-batch1-rep1 | TT-seq | new |
| FLV-rep2 | ATMP-seq | new | FLV+TOP1-IAA+ICRF-rep1 | ATMP-seq | new | TT-TOP1-IAA+ICRF-batch1-rep2 | TT-seq | new |
| TPL+FLV-rep1 | ATMP-seq | new | FLV+TOP1-IAA+ICRF-rep2 | ATMP-seq | new | TT-TOP1-IAA+ICRF-batch3-rep3 | TT-seq | new |
| TPL+FLV-rep2 | ATMP-seq | new | Prometaphase-rep1 | ATMP-seq | new | TT-RAD21-IAA-batch1-rep1 | TT-seq | new |
| nuclei-rep1 | ATMP-seq | new | Prometaphase-rep2 | ATMP-seq | new | TT-RAD21-IAA-batch1-rep2 | TT-seq | new |
| nuclei-rep2 | ATMP-seq | new | Prometaphase-Ctrl-gDNA | ATMP-seq | new | TT-CTCF-5phiAA-batch1-rep1 | TT-seq | new |
| nuclei+NTPs-rep1 | ATMP-seq | new | Asyn-Ctrl-gDNA | ATMP-seq | new | TT-CTCF-5phiAA-batch1-rep2 | TT-seq | new |
| nuclei+NTPs-rep2 | ATMP-seq | new | Prometaphase-CAPH1-IAA-rep1 | ATMP-seq | new | TT-ICRF-batch1 | TT-seq | new |
| nuclei-TOP1 | ATMP-seq | new | Prometaphase-CAPH1-IAA-rep2 | ATMP-seq | new | TT-ICRF+CPT-batch1 | TT-seq | new |
| eHAP-rep1 | ATMP-seq | new | Prometaphase-CAPH2-IAA-rep1 | ATMP-seq | new | TT-TOP1-IAA-batch1-rep1 | TT-seq | published |
| eHAP-rep2 | ATMP-seq | new | Prometaphase-CAPH2-IAA-rep2 | ATMP-seq | new | TT-TOP1-IAA-batch2-rep2 | TT-seq | new |
| ICRF+CPT-rep1 | ATMP-seq | new | TOP2A-IAA | ATMP-seq | published | TT-TOP1-IAA-batch2-rep3 | TT-seq | new |
| ICRF+CPT-rep2 | ATMP-seq | new | TOP2B-5phiAA | ATMP-seq | published | TOP2Acc-untreated | Cleavage-seq | new |
| TOP1-IAA+ICRF-rep1 | ATMP-seq | new | GM12878-ATMP-rep1 | ATMP-seq | published | TOP2Acc-TOP1-AID-IAA | Cleavage-seq | new |
| TOP1-IAA+ICRF-rep2 | ATMP-seq | new | GM12878-ATMP-rep2 | ATMP-seq | published | TOP2Bcc-untreated | Cleavage-seq | new |
| TOP2A-IAA+CPT-rep1 | ATMP-seq | new | GM12878-Ctrl | ATMP-seq | published | TOP2Bcc-TOP1-AID-IAA | Cleavage-seq | new |
| TOP2A-IAA+CPT-rep2 | ATMP-seq | new | Drosophila-ATMP-rep1 | ATMP-seq | published |  |  |  |
| TOP2B-5phiAA+CPT-rep1 | ATMP-seq | new | Drosophila-Ctrl-rep1 | ATMP-seq | published |  |  |  |
| TOP2B-5phiAA+CPT-rep2 | ATMP-seq | new | Drosophila-ATMP-rep2 | ATMP-seq | new |  |  |  |
| ICRF | ATMP-seq | new | Drosophila-Ctrl-rep2 | ATMP-seq | new |  |  |  |
| CPT | ATMP-seq | new | yeast-ATMP | ATMP-seq | new |  |  |  |
| TOP1-IAA | ATMP-seq | new | yeast-Ctrl | ATMP-seq | new |  |  |  |
| CTCF-5phiAA | ATMP-seq | new |  |  |  |  |  |  |
| RAD21-IAA | ATMP-seq | new |  |  |  |  |  |  |

**Table 2.** Additional sequencing datasets used for analyses in response to reviewers.

| Sample name | Category | New? |
| --- | --- | --- |
| HCT116-gamma-ATMP | ATMP-seq | new |
| HCT116-gamma-TPL | ATMP-seq | new |
| HCT116-gamma-Ctrl | ATMP-seq | new |
| HCT116-Ctrl-rep3 | ATMP-seq | new |
| HCT116-TOP2B-CPT-100nM | ATMP-seq | new |
| TT-HCT116-batch3-rep1 | TT-seq | new |
| TT-HCT116-batch3-rep2 | TT-seq | new |
| TT-TOP2B-5phiAA+CPT-100nM-rep1 | TT-seq | new |
| TT-TOP2B-5phiAA+CPT-100nM-rep2 | TT-seq | new |
| TT-TOP2B-5phiAA+CPT-1uM-rep1 | TT-seq | new |
| TT-TOP2B-5phiAA+CPT-1uM-rep2 | TT-seq | new |

## Results

### Transcription contributes to large-scale supercoiling across the human genome

Transcription is a major source of DNA supercoiling [5, 6, 11]. During RNA polymerase (RNAP) elongation, DNA-tracking by the polymerase generates positive supercoiling ahead of and negative supercoiling behind the elongating complex, giving rise to local twin-supercoiled domains around genes. Whether transcription influences DNA supercoiling beyond the immediate vicinity of genes, however, remains unclear.

We observed that transcription contributes not only to the formation of local twin-supercoiled domains (Figure S1a) but also to large-scale DNA supercoiling across the genome (Figures 1a-b). This result in HCT116 cells is consistent with our previous observations in a different cell line, GM12878 [5], supporting a general contribution of transcription to genome-wide topology.

**Figure 1.**
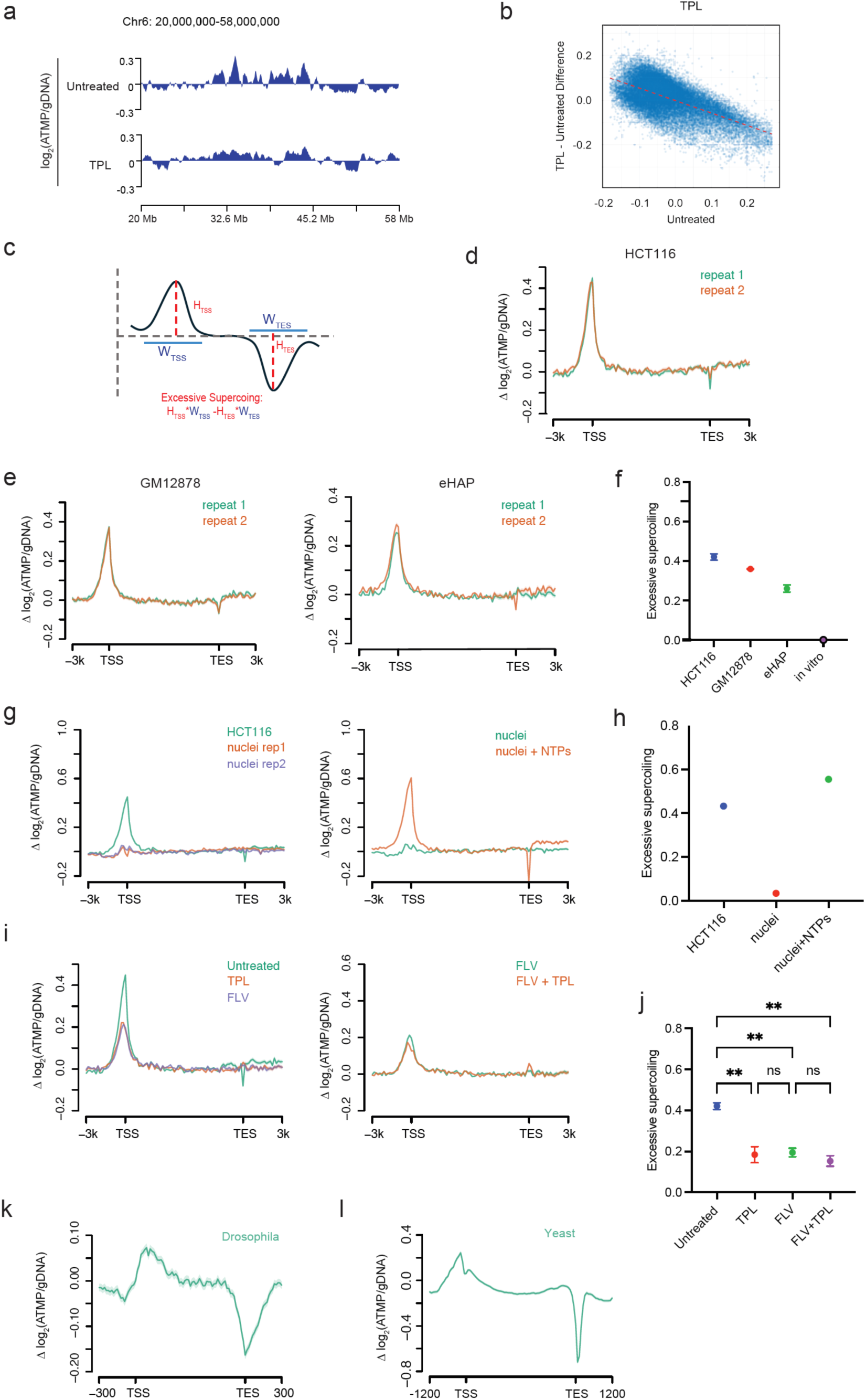
Genome-wide supercoiling and excessive negative supercoiling around genes under different transcription conditions. **a**. Example regions of genome-wide supercoiling distribution in untreated cells and upon transcription inhibition by triptolide (TPL) in HCT116 cells. Bin size: 100 kb. **b.** Changes in genome-wide supercoiling following TPL treatment in HCT116 cells. Dotted line indicates general reduction in supercoiling levels. Bin size: 100 kb. **c.** Schematic of symmetrical twin-supercoiled domains around genes in vitro and the equation for quantitative estimation of excessive negative supercoiling. **d.** Twin-supercoiled domains around genes in HCT116 cells. **e.** Twin-supercoiled domains around genes in GM12878 and eHAP cells. **f.** Quantification of excessive negative supercoiling around genes in living cells and in vitro. **g.** Supercoiling around genes in living cells, extracted nuclei, and extracted nuclei with NTP addition. **h.** Quantification of excessive negative supercoiling around genes under conditions shown in (g). **i.** Supercoiling around genes under untreated condition, transcription inhibition by TPL (blocks RNAP initiation and eliminates elongation), transcription elongation inhibition by flavopiridol (FLV), and combined TPL + FLV treatment. **j.** Quantification of excessive negative supercoiling under various transcription inhibition conditions. Significance was calculated by one-way ANOVA analysis based on two replicates per condition (*p < 0.05, **p < 0.01, ***p < 0.001, ****p < 0.0001). **k.** Twin-supercoiled domains around genes in *Drosophila* S2 cells. **l.** Twin-supercoiled domains around genes in G1-synchronized yeast cells.

However, this finding presents a conceptual paradox. Biophysical modeling [6] and in vitro measurements [14] indicate that transcription generates equal amounts of negative and positive supercoiling within twin-supercoiled domains (Figure 1c). Under this framework, equal and opposite supercoils generated locally by transcription should cancel when averaged at larger scales, yielding no net contribution to genome-wide supercoiling.

### Excessive negative supercoiling within transcription-driven twin-supercoiled domains

Interestingly, quantitative analysis of twin-supercoiled domains revealed a marked asymmetry: negative supercoiling at transcription start sites (TSS) exceeded positive supercoiling at transcription end sites (TES) (Figure 1d), resulting in a net excess of negative supercoiling around genes (Figures 1f), as inferred from normalized ATMP incorporation density [5, 15]. This imbalance contrasts with the symmetric positive and negative supercoiling generated during transcription in vitro (Figures 1c and 1f).

Excessive negative supercoiling was present in multiple human cell lines (Figures 1e-f) and was broadly observed across genes. Its magnitude scaled with transcriptional activity (Figure S1a) and correlated quasi-quantitatively with gene length and expression level (Figures S1b-c). These results suggest that asymmetric supercoiling accumulation during transcription could underlie the emergence of large-scale genome-wide supercoiling, thereby resolving the discrepancy with biophysical models.

To determine the origin of this imbalance, we isolated nuclei from living cells [16]. These nuclei retain static chromatin features, including DNA sequence, nucleosome positioning, chromatin accessibility, and higher-order chromatin architecture, but lack transcription and other ATP-dependent processes. In extracted nuclei, excessive negative supercoiling was abolished (Figures 1g-h), excluding static chromatin features or supercoiling measurement artifacts as its source. Treatment with calf thymus topoisomerase I, which relaxes both negative and positive supercoils, produced no further change (Figure S1d), confirming the absence of active chromatin processes that could generate dynamic supercoiling in the nuclei. Strikingly, reactivation of transcription by NTP supplementation restored twin-supercoiled domains and excessive negative supercoiling around genes (Figures 1g-h), demonstrating that active transcription is required to generate this asymmetry.

### RNAP elongation, rather than initiation, generates excessive negative supercoiling

To determine which stage of transcription generates excessive negative supercoiling, we compared the effects of inhibitors targeting distinct steps of transcription. Triptolide (TPL), which blocks transcription initiation and consequently eliminates elongation over time, reduced excessive negative supercoiling to a similar extent as flavopiridol (FLV), a specific inhibitor of RNAP elongation (Figures 1i-j). Combined inhibition of initiation and elongation produced no additional reduction (Figures 1i-j), indicating that RNAP elongation, rather than initiation or promoter-proximal events including the short-distance RNAP movement prior to stable elongation, is responsible for generating excessive negative supercoiling.

Promoter-proximal-paused RNAP has been reported to stimulate BAF-mediated nucleosome eviction at TSS [17], which could in principle influence local supercoiling. However, BAF inhibition did not alter supercoiling when elongation was blocked (Figure S1e). Likewise, inhibition of histone deacetylase (HDAC), which enhances nucleosome turnover [18], had no effect on excessive negative supercoiling in the absence of elongation (Figure S1e), arguing against contributions from chromatin remodeling independent of elongation.

Chromatin buffering has been reported to absorb positive supercoiling generated ahead of the elongating RNAP in vitro [19], potentially contributing to the observed excessive negative supercoiling in living cells. To examine this possibility, we performed ATMP-seq in *Drosophila* S2 cells and G1-synchronized budding yeast cells. Both are eukaryotic systems in which DNA is packed into nucleosomes and organized into chromatin and therefore, exhibit similar chromatin-mediated buffering for positive supercoiling absorption. Interestingly, although negative supercoiling at TSS remains somewhat greater than positive supercoiling at TES in both systems, the asymmetry between the twin-supercoiled domains is substantially reduced compared with that observed in human cells (Figures 1k-l). Notably, in yeast, positive supercoiling accumulates significantly at TES in a manner consistent with predictions based on gene orientation (Figure S1f). These results suggest that chromatin-mediated absorption of positive supercoiling is unlikely to be a major contributor to the excessive negative supercoiling observed around human genes.

Because divergent transcription is widespread at human promoters, we also tested whether antisense transcription contributes to excessive negative supercoiling. Although CTCF depletion increases antisense transcription at a subset of genes without affecting canonical transcription in HCT116 cells [20], auxin-induced degradation of CTCF in the same cell line did not alter excessive negative supercoiling (Figure S1g), indicating that divergent transcription is not its primary source.

Finally, we asked whether the residue negative supercoiling observed after elongation inhibition could be explained by incomplete suppression of transcription. Despite FLV treatment, low levels of transcription persisted genome-wide (Figure S1h), and genes with higher residual transcription exhibited correspondingly higher levels of residual excessive negative supercoiling (Figure S1i). This correlation supports the conclusion that the remaining supercoiling reflects lingering RNAP elongation activity, further reinforcing RNAP elongation as the source of excessive negative supercoiling around genes.

### Human topoisomerases preferentially relax positive supercoils during transcription elongation

Although RNAP elongation generates equal amounts of negative and positive supercoiling in vitro, elongation in living human cells results in a net accumulation of negative supercoiling. We hypothesized that this imbalance arises not from asymmetric supercoiling generation but from differential supercoiling relaxation. In particular, DNA topoisomerases, absent from in vitro assays but active in cells, may preferentially relax positive supercoils during transcription elongation [21, 22].

Human topoisomerase I (TOP1) is catalytically active along gene bodies and is stimulated by elongating RNAPII through interactions with its Serine 2-phoshorylated C-terminal domain (CTD) [23]. To test whether elongating RNAPII enhances TOP1 activity in a supercoiling-sensitive manner, we quantified TOP1 relaxation kinetics in the presence of Ser2-phosphorylated RNAPII CTD (RNAPII-CTD (S2P)) (Figure S2a), a hallmark of transcription elongation. RNAPII-CTD (S2P) stimulation accelerated TOP1 relaxation of positively supercoiled DNA (Figure 2a) more strongly than negatively supercoiled DNA (Figure 2b). This effect was specific to elongating form of RNAPII, as unmodified CTD did not enhance positive supercoiling relaxation (Figure 2a). Even in the absence of RNAPII stimulation, TOP1 has a modest intrinsic preference for relaxing positive supercoils (Figures S2b-c), consistent with previous reports [21].

**Figure 2.**
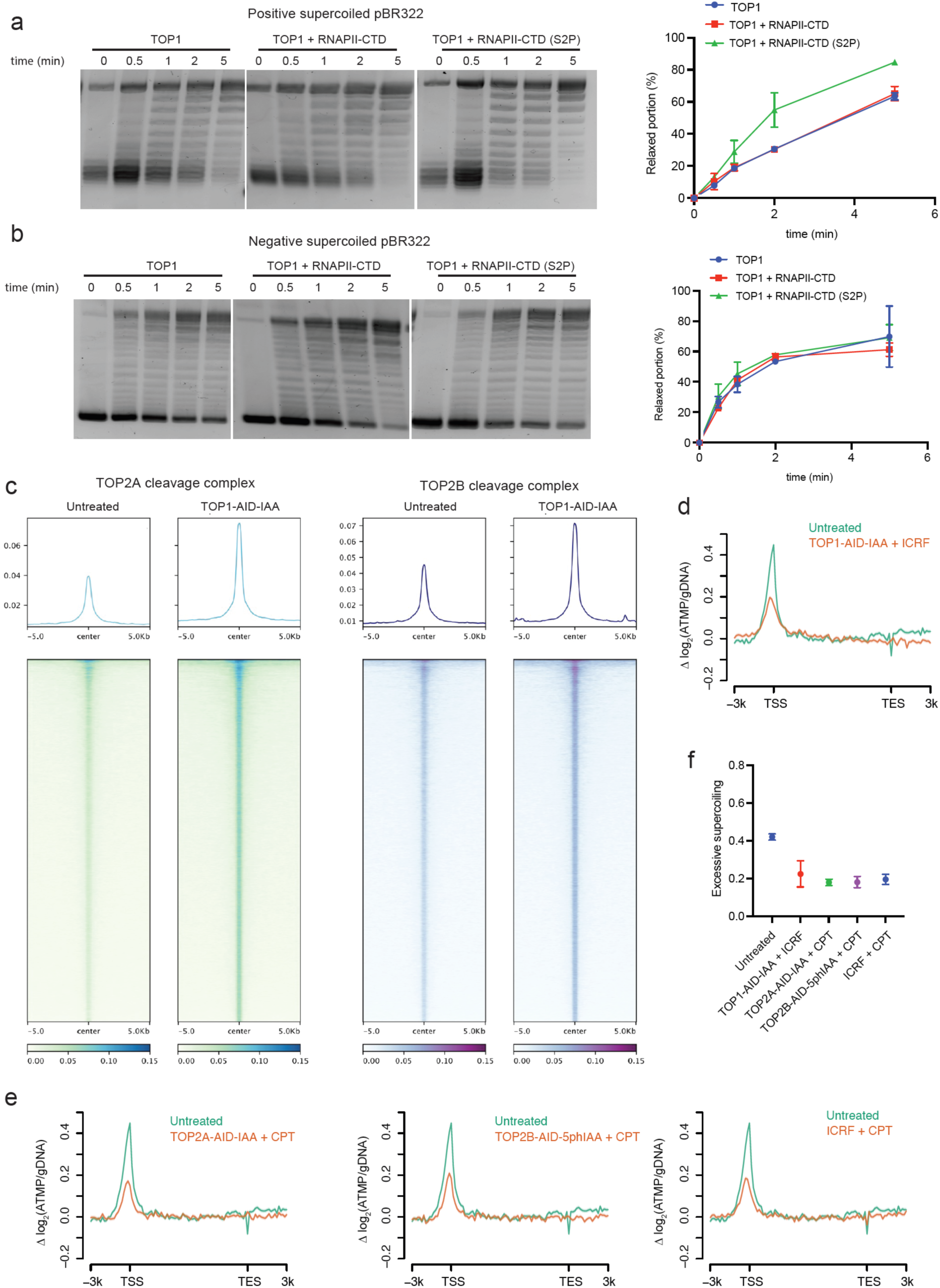
Preferential positive supercoiling relaxation during transcription elongation drives negative supercoiling around genes. **a**. Positively supercoiled pBR322 was relaxed by human TOP1, TOP1 + RNAPII-CTD, and TOP1 + RNAPII-CTD (S2P), with remaining supercoiling level at indicated time points quantified by agarose gel electrophoresis (left). Positive supercoiling relaxation rate was quantified for each condition based on the gel, with normalization to time point 0 and from two replicates per condition (right). **b.** Negatively supercoiled pBR322 was relaxed by human TOP1, TOP1 + RNAPII-CTD, and TOP1 + RNAPII-CTD (S2P), with remaining supercoiling level at indicated time points quantified by agarose gel electrophoresis (left). Negative supercoiling relaxation rate was quantified for each condition based on the gel, with normalization to time point 0 and from two replicates per condition (right). **c.** Levels of TOP2Acc and TOP2Bcc peaks across the human genome, in untreated cells and upon TOP1 AID depletion. **d.** Supercoiling around genes in untreated cells and upon simultaneous perturbation of TOP1 and TOP2 via TOP1 AID depletion and TOP2 inhibitor ICRF-193 (ICRF) treatment. **e.** Supercoiling around genes in untreated and upon simultaneous perturbation of TOP1 and TOP2 via TOP2 AID depletion and TOP1 inhibitor camptothecin (CPT) treatment, or via combined ICRF + CPT treatment. **f.** Quantification of excessive negative supercoiling under various conditions of simultaneous TOP1 and TOP2 perturbation, from two replicates per condition.

In addition, topoisomerase II (TOP2) may provide additional asymmetric relaxation. Human TOP2 has been shown to preferentially relax positive supercoils in vitro [24] and has been implicated in transcription elongation [9]. Acute depletion of TOP1 using an auxin-inducible degron (AID) system led to increased genome-wide catalytic engagement of TOP2A and TOP2B, as measured by their cleavage complexes (TOP2Acc and TOP2Bcc) (Figure 2c). These findings are consistent with physical interaction and functional compensation between topoisomerases [25–27]. Moreover, compared with naked DNA templates, chromatin may further enhance topoisomerase activity for preferential relaxation of positive supercoiling over negative supercoiling [28, 29]. Together, these results provide a mechanistic basis for the preferential removal of positive supercoils by both TOP1 and TOP2 during RNAP elongation in human cells.

### Preferential relaxation of positive supercoils during RNAP elongation drives negative supercoiling around genes

To test whether negative supercoiling around genes arises from preferential relaxation of positive supercoils during RNAP elongation, we examined supercoiling profiles following topoisomerase perturbation.

Acute AID depletion of TOP1 did not reduce negative supercoiling (Figure S2d). Instead, supercoiling increased modestly at both TSS and TES (Figures S2d and S2f), consistent with the established role of TOP1 in global supercoiling relaxation across the genome [5]. Depletion of either TOP2A or TOP2B produced similar moderate increase in supercoiling (Figures S2e-f), indicating that both isoforms contribute comparably to global supercoiling relaxation. Likewise, pharmacological inhibition TOP1 or TOP2 alone did not significantly reduce negative supercoiling (Figures S2e-f). Together, these results indicate that perturbation of either enzyme individually is insufficient to disrupt the asymmetric supercoiling relaxation during RNAP elongation, consistent with functional redundancy between TOP1 and TOP2.

We therefore examined the consequence of simultaneous disruption of TOP1 and TOP2. Co-perturbation of both enzymes led to a significant reduction in negative supercoiling around genes, despite a predicted loss of global supercoiling relaxation (Figures 2d and 2f). This reduction was observed across multiple dual TOP1/TOP2 perturbation strategies (Figures 2e-f) and could not be attributed to major decrease in transcriptional activity (Figure S2g), supporting a model in which preferential relaxation of positive supercoils by human topoisomerases during RNAP elongation drives the accumulation of negative supercoiling around genes.

### Transcription-driven negative supercoiling contributes to large-scale genome-wide supercoiling

We hypothesized that transcription-driven negative supercoiling extends beyond individual genes and thereby contributes to large-scale genome-wide supercoiling.

Consistent with this idea, highly transcribed genomic regions exhibited substantial accumulation of negative supercoiling (Figure 3a). We next examined gene-rich topologically associating domain (TAD) boundaries, which are frequently enriched for transcriptional activity. As previously reported [5], we confirmed negative supercoiling accumulation at TAD boundaries (Figure 3b). This accumulation was abolished in extracted nuclei lacking transcriptional activity and restored upon transcription reactivation (Figure 3c), excluding a role for static chromatin features.

**Figure 3.**
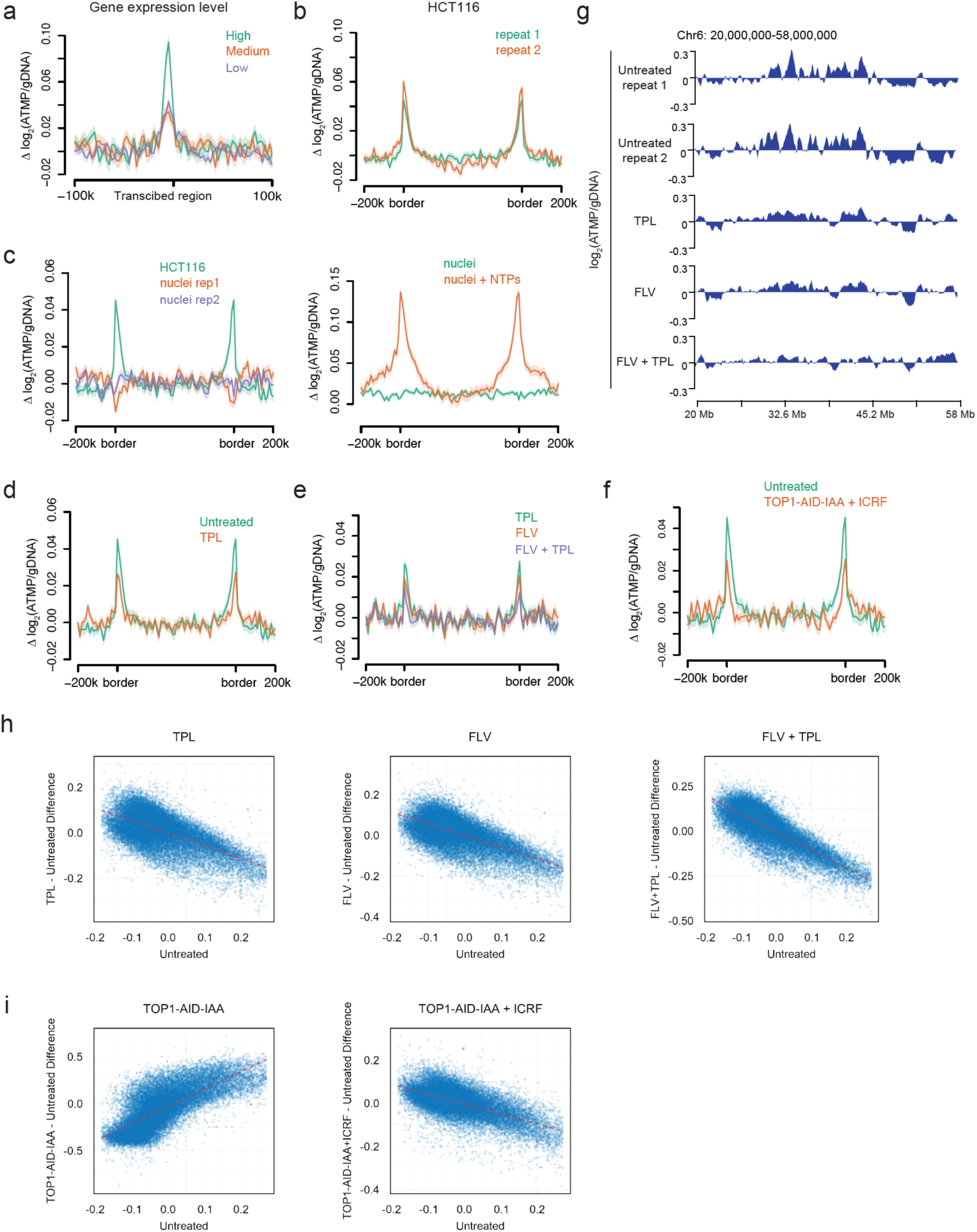
Transcription-driven negative supercoiling contributes to large-scale genome-wide supercoiling. **a**. Supercoiling around genomic regions with transcription enrichment. Gene-rich TAD boundary regions are categorized into three groups based on the overall gene expression level across the region. **b.** Supercoiling at TAD boundaries in living cells. **c.** Supercoiling at TAD boundaries in living cells, extracted nuclei, and extracted nuclei with NTP addition. **d.** Supercoiling at TAD boundaries in untreated cells and upon transcription inhibition via TPL treatment. **e.** Supercoiling at TAD boundaries upon treatments of TPL, FLV, and TPL + FLV. **f.** Supercoiling at TAD boundaries in untreated cells and upon simultaneous TOP1 and TOP2 perturbation. **g.** Example regions of genome-wide supercoiling distribution in untreated cells and under various transcription inhibition conditions. Bin size: 100 kb. **h.** Changes in genome-wide supercoiling under various transcription inhibition and topoisomerase perturbation conditions. Dotted lines indicate general reduction or elevation in supercoiling levels. Bin size: 100 kb.

Negative supercoiling at TAD boundaries required active transcription (Figure 3d) and specifically RNAP elongation rather than initiation (Figure 3e). Individual perturbation of TOP1 and TOP2 further increased negative supercoiling at TAD boundaries (Figure S3a), whereas simultaneous disruption of both enzymes markedly reduced it (Figures 3f and S3b), mirroring the behavior observed around individual genes. These results indicate that transcription-driven negative supercoiling extends across larger genomic regions.

To assess whether this effect extends beyond the hundred-kb TAD boundaries, we examined megabase-scale supercoiling. Strikingly, genome-wide supercoiling at this scale similarly dependent on RNAP elongation but not initiation (Figures 3g-h), and displayed comparable sensitivity to topoisomerase perturbations (Figures 3i and S3c-d). Together, these findings demonstrate that transcription-driven negative supercoiling is a major contributor to large-scale genome-wide topology in human cells.

### Cohesin-mediated loop extrusion contributes to genome-wide supercoiling

Genome-wide supercoiling exhibited a moderate correlation with transcriptional activity on the large scale (Figure S4a), consistent with transcription as a major contributor to genome-wide topology. However, the imperfect correlation suggested the existence of additional mechanisms shaping supercoiling across the human genome.

We first excluded large-scale regional differences in topoisomerase relaxation activity as a primary driver. Although topoisomerases preferentially relax transcription-generated supercoils locally around genes, perturbation experiments indicated that at large genomic scales, topoisomerase-mediated relaxation occurs relatively uniformly without strong regional bias (Figures S4b-c).

We also excluded major contributions from static chromatin features, including DNA sequence effects on looping efficiency [30] and nucleosome-based determinants of 3D chromatin compartments [31]. Comparing supercoiling measurements normalized to genomic DNA and to extracted nuclei revealed minimal influence of static chromatin features on large-scale genome-wide topology (Figures S4d-e).

Interestingly, transcription reactivation in extracted nuclei largely restored genome-wide supercoiling patterns characteristic of living cells (Figures S4f-g). The increased magnitude of supercoiling likely reflects elevated transcription following NTP addition (Figures S4f-g). These observations indicate that ATP-dependent chromatin processes, in addition to transcription itself, contribute to large-scale supercoiling.

Among potential candidates, cohesin-mediated loop extrusion is an ATP-dependent chromatin process that operates genome-wide and has been shown in vitro to generate negative supercoiling in extruded DNA [12]. We therefore examined its role in shaping genome-wide supercoiling.

Acute AID-mediated cohesin depletion altered supercoiling distribution patterns (Figures 4a-b), indicating that cohesin contributes to genome-wide supercoiling in living cells. Conversely, AID depletion of CTCF, which enhances cohesin loop extrusion by removing extrusion barriers, produced distinct changes in genome-wide supercoiling distribution (Figures 4a-b). These effects were not explained by changes in transcriptional activity (Figures S5a-b), supporting cohesin-mediated loop extrusion as an independent factor contributing to large-scale genome-wide supercoiling.

**Figure 4.**
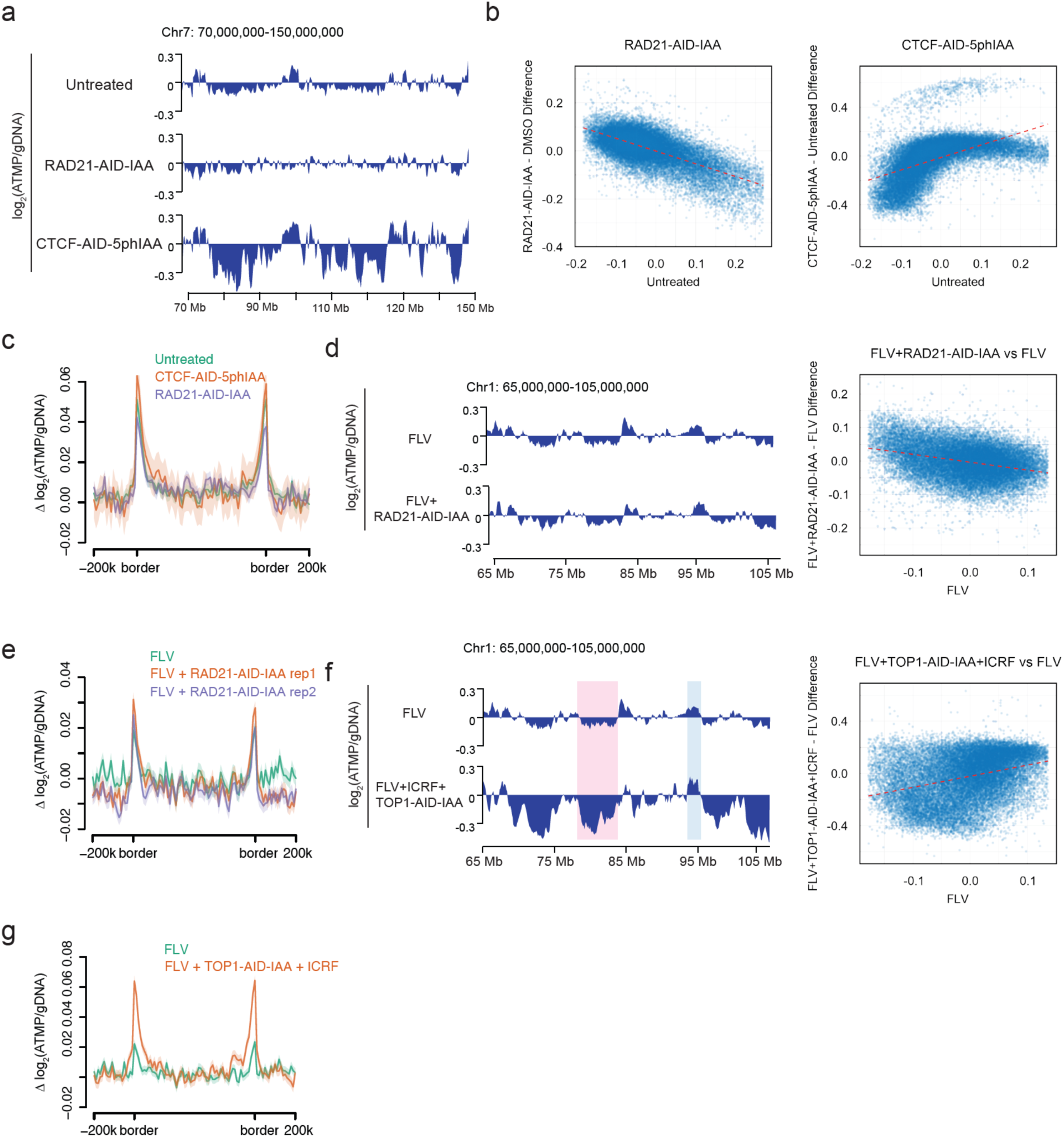
Cohesin-mediated loop extrusion contributes to genome-wide supercoiling. **a**. Example regions of genome-wide supercoiling distribution in untreated cells, upon cohesin AID depletion, and CTCF AID depletion. Bin size: 100 kb. **b.** Changes in genome-wide supercoiling upon cohesin AID depletion and CTCF AID depletion. Dotted lines indicate general reduction or elevation in supercoiling levels. Bin size: 100 kb. **c.** Supercoiling at TAD boundaries upon cohesin AID depletion and CTCF AID depletion. **d.** Example regions of genome-wide supercoiling distribution upon indicated perturbations (left), and changes in genome-wide supercoiling upon cohesin AID depletion under transcription inhibition condition. Dotted lines indicate general reduction in supercoiling levels. Bin size: 100 kb (right). **e.** Supercoiling at TAD boundaries upon indicated perturbations. **f.** Example regions of genome-wide supercoiling distribution upon indicated perturbations (left), and changes in genome-wide supercoiling upon TOP1 and TOP2 perturbation under transcription inhibition condition. Dotted lines indicate general elevation in supercoiling levels. Bin size: 100 kb (right). **g.** Supercoiling at TAD boundaries upon indicated perturbations.

### Topoisomerases relax cohesin-generated supercoiling at TAD boundaries

Although cohesin contributes to genome-wide supercoiling, depletion of cohesin or CTCF produced only minor changes in supercoiling at TAD boundaries, where cohesin frequently stalls at CTCF-binding sites (Figure 4c), consistent with previous observations [5]. Given the enrichment of topoisomerases [32] and TOPcc-mediated DNA breaks at TAD boundaries [33, 34], we hypothesized that cohesin-generated supercoiling at these sites is rapidly relaxed by topoisomerases.

To isolate cohesin-dependent effects from transcription-driven supercoiling, we performed perturbations under transcription-inhibited conditions. Because transcription inhibition could potentially influence cohesin activity, we first verified that, in the absence of transcription, cohesin depletion continued to alter large-scale supercoiling (Figure 4d) but did not affect supercoiling at TAD boundaries (Figure 4e), mirroring results obtained under transcription-active conditions.

Under transcription-inhibited conditions, we found that topoisomerase perturbation caused a global alteration in genome-wide supercoiling (Figure 4f), indicating that topoisomerases broadly relax cohesin-generated supercoiling, potentially with varying efficiencies across the genome (Figure 4f). Notably, topoisomerase disruption led to increased negative supercoiling specifically at TAD boundaries (Figure 4g), suggesting topoisomerase-mediated rapid local supercoiling relaxation at these sites under normal conditions. This effect is unlikely to reflect residual transcription, as transcription-driven negative supercoiling would be expected to decrease upon topoisomerase perturbation. However, we cannot fully exclude the possibility that topoisomerase disruption indirectly influences cohesin loading or extrusion dynamics, particularly given that loop extrusion may itself be sensitive to DNA topology [35].

### Condensin establishes an overall positively supercoiled mitotic genome

Because all SMC complexes generate negative supercoiling in extruded DNA in vitro [13], we examined whether condensin shapes genome-wide supercoiling during mitosis, when condensin, rather than cohesin, is the dominant loop extruder. Using AID depletion of condensin subunits (Figure S6a) in nocodazole-synchronized prometaphase cells (Figure S6c), we found that both condensin I and II contribute to genome-wide supercoiling during mitosis (Figure 5a). These effects are unlikely to arise from alterations in transcription or cohesin-mediated loop extrusion, as both processes are strongly repressed during mitosis [36–38].

**Figure 5.**
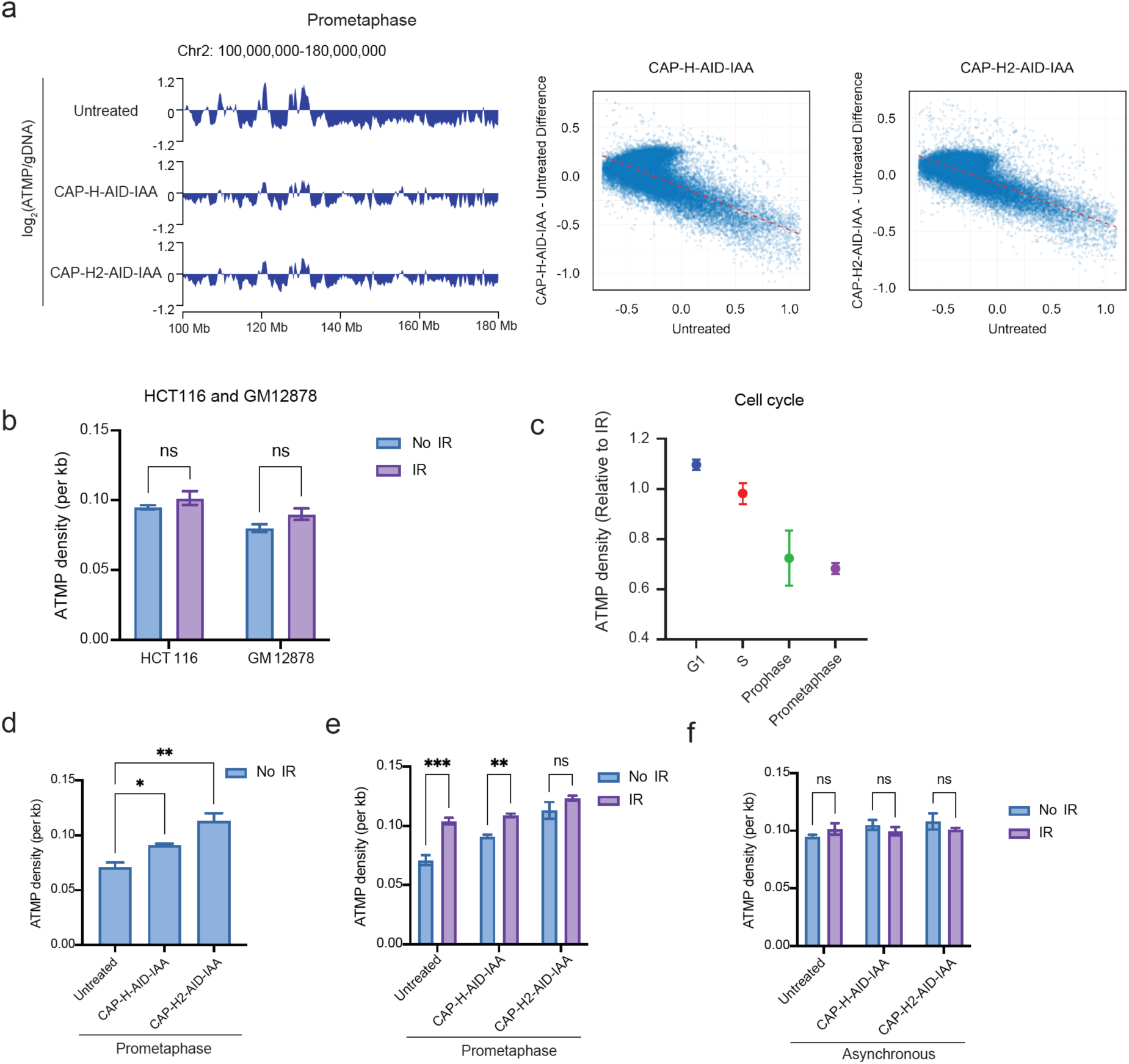
Condensin establishes an overall positively supercoiled mitotic genome. **a**. Example regions of genome-wide supercoiling distribution in prometaphase cells under conditions of untreated, condensin I subunit CAP-H AID depletion, and condensin II subunit CAP-H2 AID depletion (left). Changes in genome-wide supercoiling under condensin AID depletion conditions. Dotted lines indicate general reduction in supercoiling levels. Bin size: 100 kb (right). **b.** Average supercoiling level across the genome in HCT116 cells and GM12878 cells. ATMP density is determined by the ratio between pull-down ATMP-adducted DNA and input DNA, and compared between living cells with DNA breaks (IR: gamma irradiation) and with intact genome (No IR). Significance was calculated by two-way ANOVA analysis based on two replicates per condition (*p < 0.05, **p < 0.01, ***p < 0.001, ****p < 0.0001). **c.** Average supercoiling level across the genome in HCT116 cells at different cell-cycle stages. ATMP density is determined by the ratio between pull-down ATMP-adducted DNA and input DNA. ATMP density relative to IR is plotted by comparing ATMP density in living cells (No IR) to that in cells receiving gamma irradiation to introduce DNA breaks. **d.** Average supercoiling level across the genome in HCT116 cells at prometaphase, under conditions of untreated, condensin I AID depletion, and condensin II AID depletion. Significance was calculated by one-way ANOVA analysis based on two replicates per condition (*p < 0.05, **p < 0.01, ***p < 0.001, ****p < 0.0001). **e.** Average supercoiling level across the genome in HCT116 cells at prometaphase, with and without gamma irradiation, under conditions of untreated, condensin I AID depletion, and condensin II AID depletion. Significance was calculated by two-way ANOVA analysis based on two replicates per condition (*p < 0.05, **p < 0.01, ***p < 0.001, ****p < 0.0001). **f.** Average supercoiling level across the genome in asynchronous HCT116 cells, with and without gamma irradiation, under conditions of untreated, condensin I AID depletion, and condensin II AID depletion. Significance was calculated by two-way ANOVA analysis based on two replicates per condition (*p < 0.05, **p < 0.01, ***p < 0.001, ****p < 0.0001).

Furthermore, unlike condensin’s loop extrusion activity, its ability to constrain supercoils depends on mitosis-specific Cdk1 phosphorylation [39]. In vitro studies indicate that densely DNA-bound condensin can constrain positive supercoils in the presence of ATP [40, 41], while lower density condensin binding may instead constrain negative supercoils [42]. These findings raise the possibility that condensin modulates compensatory unconstrained supercoiling in mitotic chromosomes.

To directly assess condensin’s impact on the overall topological state of the genome, we measured the average level of genome-wide supercoiling across cell-cycle stages. G1– and S-phase populations were isolated by flow cytometry (Figure S6b), and prophase and prometaphase cells were obtained using RO-3306 and nocodazole synchronization, respectively (Figure S6c). As a relaxed reference state, we introduced widespread DNA breaks by gamma radiation [43], which efficiently relaxes supercoiling genome-wide (Figure S6d) and avoids the open chromatin region bias associated with bleomycin treatment [43].

Asynchronous, G1-, and S-phase cells exhibited a supercoiling-relaxed genome on average (Figures 5b-c). In contrast, prophase and prometaphase cells displayed a shift toward an overall positively supercoiled genome (Figure 5c), revealing a mitosis-specific global change in genome-wide DNA topology. Consistently, depletion of condensin I or II in prometaphase cells reduced this overall positive supercoiling (Figure 5d), with condensin II depletion nearly restoring a supercoiling-relaxed genome (Figure 5e). As a control, condensin depletion in interphase cells did not alter the average genome-wide supercoiling level (Figure 5f), underscoring the mitosis-specific role of condensin in shaping an overall positively supercoiled genome. Together, these results identify condensin as a central player in establishing and maintaining the overall positively supercoiled state of mitotic chromosomes.

### Negative supercoiling around genes represses transcription

To assess the functional consequences of genome-wide DNA supercoiling, we examined whether locally accumulated negative supercoiling acts as an active regulatory layer in human transcription. Genome-wide transcriptional activity was measured using transient transcriptome sequencing (TT-seq) following metabolic labeling of nascent RNA. As expected, TT-seq detected widespread antisense transcription upstream of the TSS (Figure S7a), and transcription inhibition markedly reduced nascent RNA levels genome-wide (Figure S7a).

Previous studies have suggested that negative supercoiling generally facilitates transcription [44–47], although inhibitory effects have been reported under certain conditions [47–49]. However, these conclusions were primarily derived from in vitro systems [47], exogenous reporter constructs [46, 49], or selected genes in lower organisms [45, 48]. The genome-wide transcriptional response of endogenous human genes responding to topological dynamics has remained unclear.

Strikingly, simultaneous perturbation of TOP1 and TOP2, conditions that reduce the accumulation of negative supercoiling around genes, increased transcriptional output genome-wide (Figure 6a). The effect was observed across genes spanning a broad range of expression levels (Figure 6b) and was global in nature, with only a minority of genes deviating from the overall trend (Figure 6a). Increased nascent RNA signal was most pronounced at gene 5’ ends and extended throughout gene bodies (Figure 6b), indicating enhanced transcription initiation. These findings suggest that locally accumulated negative supercoiling exerts a repressive effect on transcription in human cells.

**Figure 6.**
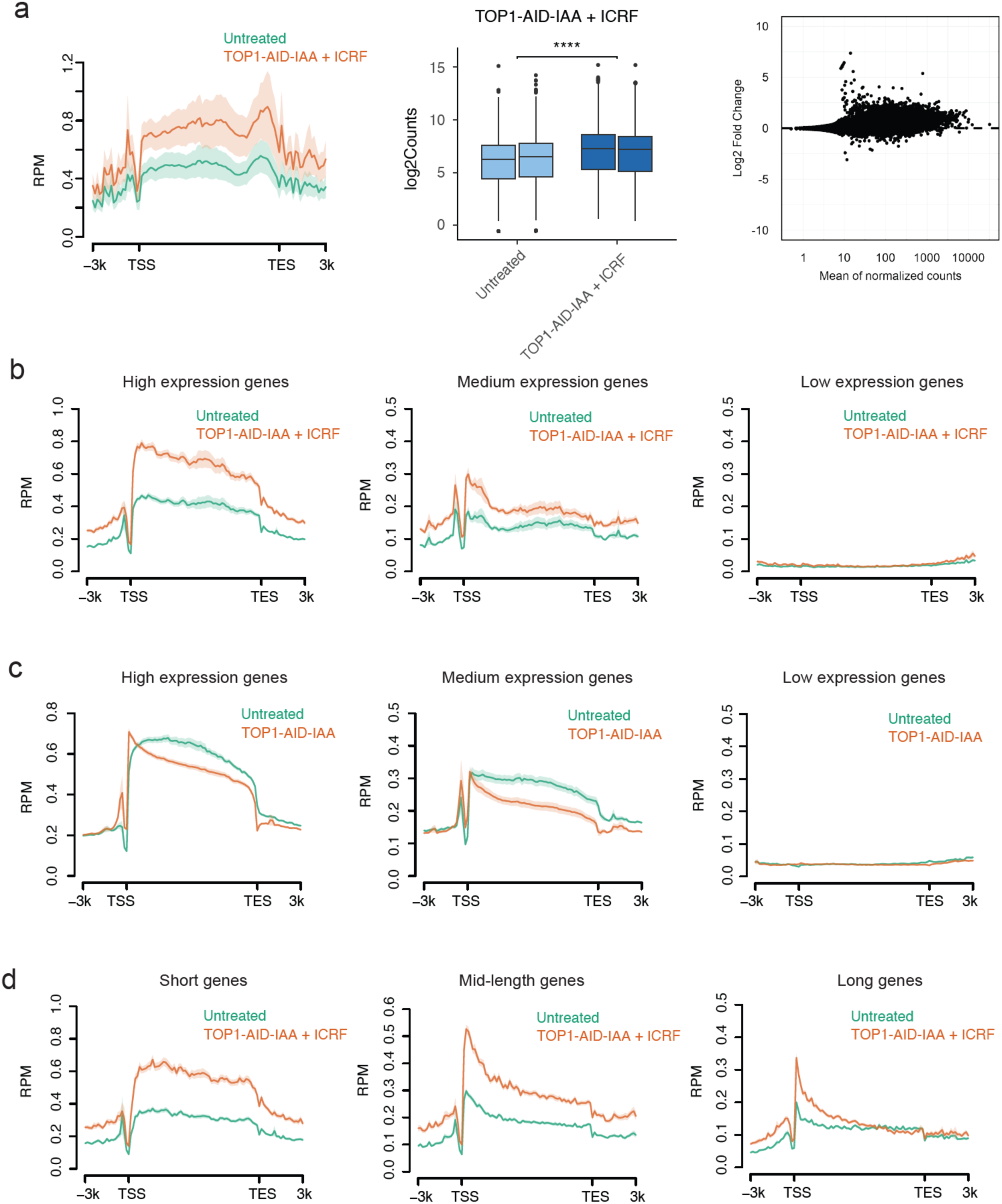
Negative supercoiling around genes represses transcription. **a**. Transcription activity by nascent RNA sequencing reads per million, compared between conditions of untreated and simultaneous TOP1 AID depletion and TOP2 ICRF inhibition, across the gene body (left), biological duplicates (middle), and individual genes (right). **b.** Transcription activity across the gene body, compared between conditions of untreated and simultaneous TOP1 AID depletion and TOP2 ICRF inhibition, for genes categories grouped by gene expression level. High expression genes are top 20%, low expression genes are bottom 50%, and medium expression genes are in between. **c.** Transcription activity across the gene body, compared between conditions of untreated and TOP1 AID depletion, for genes categories grouped by gene expression level. High expression genes are top 20%, low expression genes are bottom 50%, and medium expression genes are in between. **d.** Transcription activity across the gene body, compared between conditions of untreated and simultaneous TOP1 AID depletion and TOP2 ICRF inhibition, for gene categories grouped by gene length. Long genes are top 20%, short genes are bottom 50%, and mid-length genes are in between.

Consistently, depletion of TOP1 alone, which modestly increased negative supercoiling at the TSS (Figure S2d), did not lead to an increase in transcription across genes (Figure 6c). Likewise, brief treatment of ICRF, which did not alter negative supercoiling at the TSS (Figure S2e), produced no detectable change in transcription activity (Figure S7b). Together, these results support a model in which transcription-driven negative supercoiling acts as a constraining feedback mechanism that limits transcriptional output.

Collectively, we concluded that genome-wide DNA supercoiling generated by transcription and SMC complexes constitutes an active layer in human genome regulation of transcription.

### Topoisomerase perturbation impairs transcription elongation

We found that disruption of topoisomerase activity impairs transcription elongation. Simultaneous perturbation of TOP1 and TOP2 led to elongation defects in a gene length-dependent manner, with the strongest reductions observed in long genes (Figure 6d). Consequently, expression of long genes was disproportionately reduced. These findings align with longstanding predictions based on the established role of topoisomerases in facilitating RNAP elongation [23, 50]. They are also consistent with prior reports linking topoisomerase dysfunction to impaired long-gene expression in neurons associated with neurological disorders [51–53] and during B-cell development, where compromised long-gene transcription contributes to immunodeficiency [54, 55].

Together, these results indicate that topoisomerases play dual roles in transcription: while preferential relaxation of positive supercoils during elongation promotes accumulation of inhibitory negative supercoiling, topoisomerases are simultaneously required to relieve torsional stress that would otherwise impede RNAP elongation, particularly across long genes.

## Discussion

We demonstrated that transcription and SMC complexes together establish genome-wide DNA supercoiling in human cells (Figure 7). Transcription-driven negative supercoiling, arising from preferential relaxation of positive supercoils during elongation, represents a major source of large-scale genome-wide topology. In parallel, SMC complexes, including cohesin in interphase and condensin in mitosis, also shape genome-wide supercoiling. Functionally, the accumulation of negative supercoiling around genes represses transcription, establishing DNA topology as an active layer of genome regulation that constrains gene expression (Figure 7).

**Figure 7.**
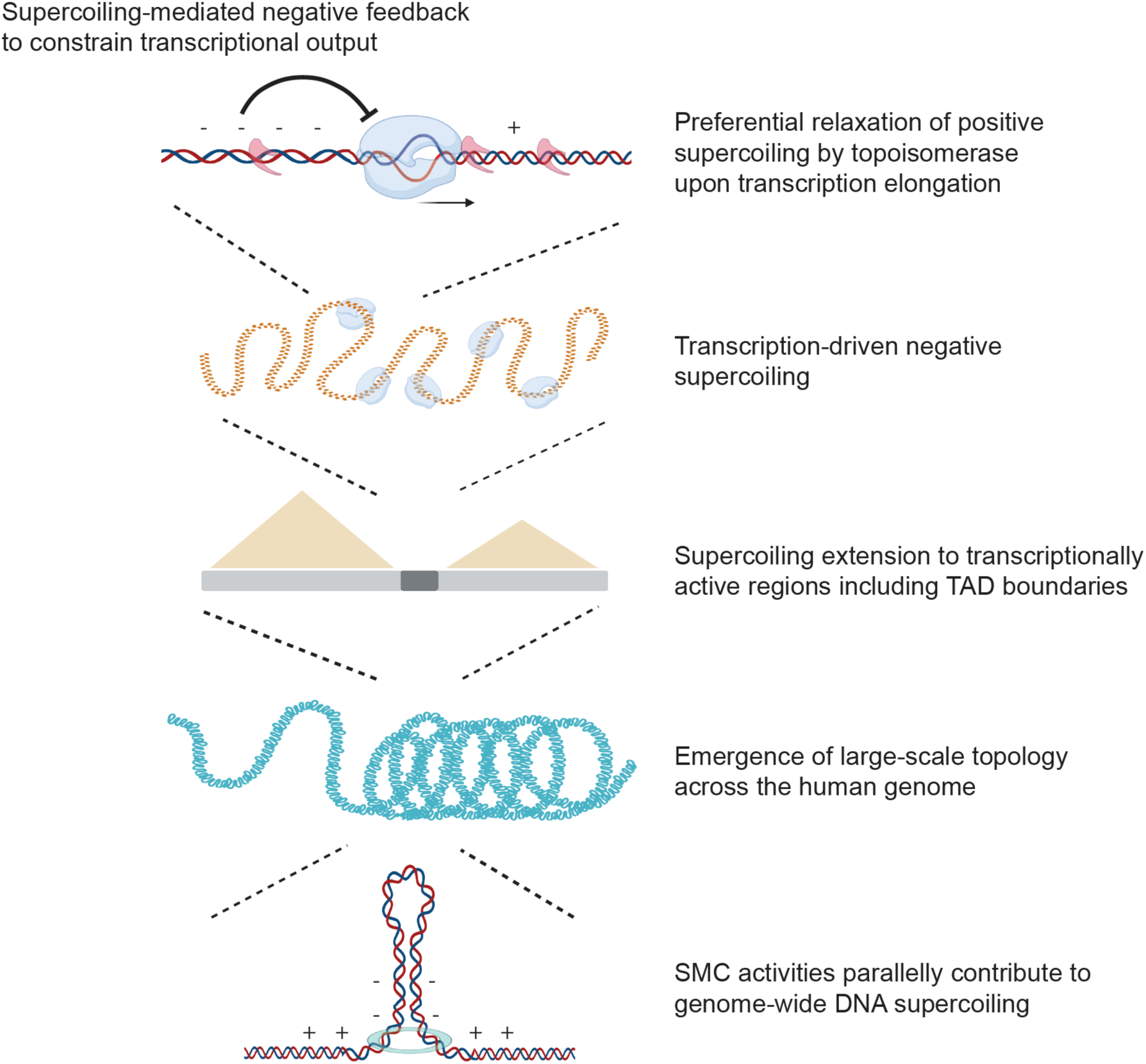
Model for the establishment and function of genome-wide DNA supercoiling in human cells. During transcription elongation, RNAP generates twin-supercoiled domains, with positive supercoils ahead of and negative supercoils behind the elongating complex. Human topoisomerases preferentially relax positive supercoils, resulting in a net accumulation of negative supercoiling around genes. This imbalance of supercoiling extends beyond individual genes to transcriptionally active regions including TAD boundaries and large-scale topology across the human genome. SMC activities, including cohesin-mediated loop extrusion in interphase and condensin in mitosis, parallelly contribute to genome-scale DNA supercoiling. The accumulation of negative supercoiling imposes a supercoiling-mediated negative feedback that constrains transcriptional output in human cells.

Although transcription and SMC complexes are principal drivers of genome-wide supercoiling, additional chromatin processes may contribute. Beyond the replicative helicase during DNA replication as a major source in proliferating cells [56], other ATP-dependent chromatin dynamics could influence local topology. Protein binding and unbinding events, including nucleosome turnover, can alter the balance between constrained and unconstrained supercoils [57]. While the amount of supercoiling generated by nucleosome turnover is typically smaller than that produced by DNA-tracking processes (Figure S8a), it may modulate local topology. Indeed, under transcription-inhibited conditions (Figure S8b), HDAC-inhibitor-induced enhancement of nucleosome turnover increased negative supercoiling, particularly within accessible chromatin regions (Figures S8c-d). However, indirect effects on cohesin dynamics cannot be excluded.

Further studies should investigate coordination between topoisomerases, transcription, and SMC complexes. Interactions between elongating RNAP and TOP2A/B may parallel those established for TOP1. The interplay between SMC complexes and topoisomerases likely vary across genomic contexts and may influence local and large-scale supercoiling across the genome. Because transcription and SMC activity both generate and respond to DNA supercoiling [35, 58–60], their effects may be additive or antagonistic depending on chromatin context and topoisomerase activity, positioning topoisomerases as central mediators of genome-wide topological homeostasis [1, 35].

Functionally, we show that negative supercoiling constrains, rather than promotes, transcription in human cells. The molecular basis of this inhibition remains to be defined. Negative supercoiling may influence multiple steps of transcriptional regulation, including transcription factor binding, chromatin remodeling, RNAP recruitment, promoter melting, and promoter clearance. In contrast, elongation defects observed upon topoisomerase disruption likely reflect transient supercoiling stress generated during RNAP progression that is not fully captured by bulk measurements of average DNA supercoiling. Together, these observations suggest dual roles of topology-based transcription regulation, that topoisomerase activities facilitate elongation yet also accumulate negative supercoiling as a transcriptional constraint.

Beyond transcription, genome-wide supercoiling may influence additional chromatin processes[61, 62]. Negative supercoiling could impede replication fork progression [63], while positive supercoiling may facilitate sister-chromatid decatenation during mitosis [64]. Exploring supercoiling and topoisomerase function during DNA replication [56] and mitotic chromosome condensation [39, 65] will further clarify their broader roles.

Overall, our findings resolve the molecular basis of genome-scale supercoiling and establish DNA topology as a dynamic regulatory feature of the human genome. The modulation of genome-wide topological homeostasis by transcription, SMC complexes, and topoisomerases may represent a unifying principle linking chromatin structure and genome function.

It is worth noting that ATMP-seq captures a population-averaged snapshot of genome-wide DNA supercoiling. At any given time, a gene may be inactive in some cells and transcribed by RNAPs at different positions along the gene body in other cells. Because an elongating RNAP generates negative supercoiling upstream and positive supercoiling downstream, these signals largely average out across gene bodies at the population level. In contrast, supercoiling at gene boundaries does not fully cancel, resulting in negative supercoiling enrichment at TSS and positive supercoiling enrichment at TES detected by ATMP-seq.

The population-averaged nature of ATMP-seq is also important for interpreting long-range supercoiling. Our model suggests that transcription-generated supercoiling can propagate beyond genes into surrounding genomic regions, although such long-range propagation is neither instantaneous nor unconstrained and may be impeded or slowed by chromatin features. Because transcription is transient and stochastic in individual cells, long-range propagation does not necessarily produce large domains of uniform supercoiling in population-averaged ATMP-seq profiles.

In addition, ATMP-seq measures DNA supercoiling levels relative to local and genome-wide baseline rather than the absolute supercoiling levels. Thus, an apparent increase in positive supercoiling in ATMP-seq measurements may reflect either increased absolute positive supercoiling or reduced negative supercoiling.

## Limitations

We employed both drug inhibition and AID-mediated depletion as acute perturbation strategies to cross-validate our findings, recognizing that each method has inherent limitations. Drug treatments act rapidly but may introduce off-target effects, whereas AID-mediated depletion, although rapid upon induction, can exhibit auxin-independent “leakiness”, causing partial protein reduction prior to induction [66]. For example, while comparisons between topoisomerase-depleted and wild-type cells indicated comparable contributions of TOP2A and TOP2B to genome-wide supercoiling relaxation, earlier analyses using uninduced AID-tagged controls suggested a more prominent role for TOP2B [5]. This discrepancy may reflect degron leakiness in uninduced samples. Newer degradation systems, such as AID2 [67] and dTAG [68], may offer advantages in future studies.

Additionally, most experiments were performed in asynchronous populations, as genome-wide supercoiling profiles were similar between asynchronous and G1-sorted cells. Nevertheless, topoisomerase perturbation could subtly alter cell-cycle composition. Although short-term treatments are unlikely to induce major shifts, cells at different stages may respond differently to perturbation. For instance, the contribution of TOP2A to genome-wide supercoiling relaxation may primarily reflect its activity in S-and G2-phase cells, where it is most abundant. Future analyses using synchronized populations may help further refine stage-specific contributions.

## Material and Data Availability

All genome-wide DNA supercoiling and topoisomerase catalytic engagement sequencing data were deposited to GSE312079. Transient transcriptome sequencing data were deposited to GSE312084. Previously published data used in this study can be referred in Sequence Read Archive under the accession number PRJNA1003820: SRX21313612 (ATMP-seq-GM12878), SRX21313613 (ATMP-seq-GM12878-Ctrl), SRX21313610 (ATMP-seq-Drosophila), SRX21313611(ATMP-seq-Drosophila-Ctrl), SRX21313592 (ATMP-seq-TOP2A-AID-IAA), SRX22776329 (ATMP-seq-TOP2B-AID2-5phIAA), and SRX22776322 (TT-seq-TOP1-AID-IAA).

## Author Contributions

C.C. designed the overall study and wrote the manuscript. L.Z. performed most of the experiments and data analysis. Q.Y. helped with the data analysis and data organization. C.V. performed prometaphase synchronization and validated CAP-H/CAP-H2 degron system.

## Acknowledgements

We thank Shuai Li (NIH) for help in topoisomerase catalytic engagement detection and analysis, Yaqiang Cao (NIH) for help in data analysis, Christophe E. Redon (NIH) for help with gamma irradiation, Ferenc Livak and Raghad Almofeez for help with flow cytometry (NIH), Mardo Koivomagi (NIH) for CycT1-cdk9 kinase, Laura Baranello (Karolinska Institute) for TOP1-AID cell line, Christian Friberg Nielsen (Copenhagen University) in Damien Hudson lab for TOP2A-AID cell line, Rafael Casellas (MD Anderson Cancer Center) for RAD21-AID cell line, Masato Kanemaki (NIG, Japan) for CTCF-AID2 cell line, Kazuhiro Maeshima (NIG, Japan) for CAP-H-AID and CAP-H2-AID cell lines, Elissa Lei (NIH) for *Drosophila* S2 cell line, Sergei Mirkin (Tufts University) for CH1585 yeast strain, and Jun Lyu (NIH) for discussions. This study used NIH Biowulf Linux cluster.

## Supplementary Figures 1-8

**Supplementary Figure 1.**
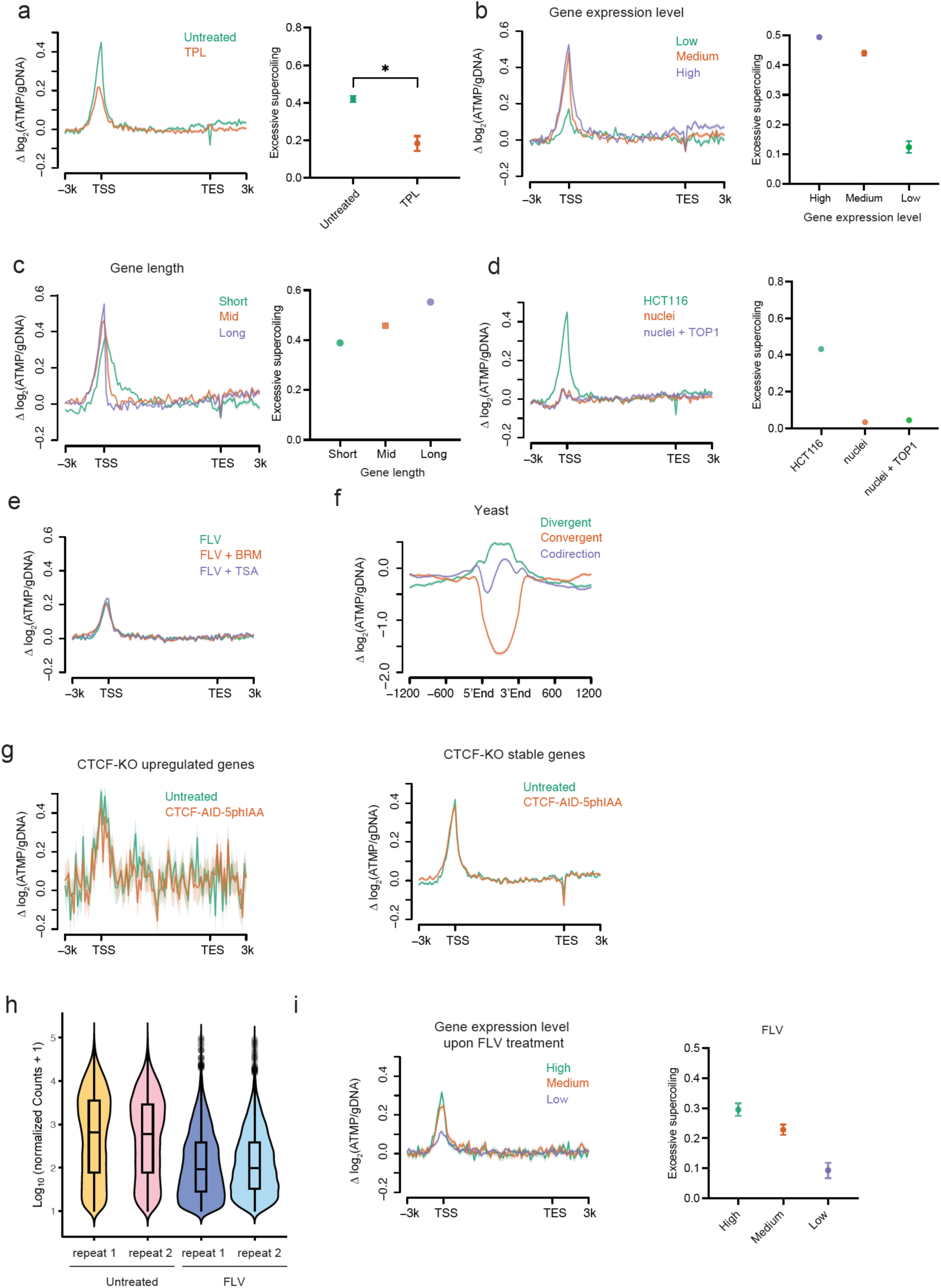
Twin-supercoiled domains under different transcription conditions. **a.** Supercoiling around genes under untreated condition and upon transcription inhibition by TPL in HCT116 cells (left), with quantification of excessive negative supercoiling (right). Significance was calculated by t-test based on two replicates per condition (*p < 0.05). **b.** Supercoiling around genes for gene categories grouped by gene expression level in HCT116 cells (left), with quantification of excessive negative supercoiling from two replicates (right). High expression genes are top 20%, low expression genes are bottom 50%, and medium expression genes are in between. **c.** Supercoiling around genes for gene categories grouped by gene length in HCT116 cells (left), with quantification of excessive negative supercoiling from two replicates (right). Long genes are top 20%, short genes are bottom 50%, and mid genes are in between. **d.** Supercoiling around genes in living cells, extracted nuclei, and extracted nuclei treated with calf thymus topoisomerase I to relax possible residue negative and positive DNA supercoiling (left), with quantification of excessive negative supercoiling (right). **e.** Supercoiling around genes upon FLV treatment, combined FLV + BRM014 (BRM) treatment, and combined FLV + trichostatin A (TSA) treatment. BRM014 is an inhibitor of BAF-mediated chromatin remodeling, and TSA is a histone deacetylase inhibitor. **f.** Supercoiling in yeast around genes with divergent, convergent, and tandem orientations. **g.** Supercoiling around genes in untreated cells and upon CTCF AID depletion in HCT116 cells, for the subset of genes with upregulated divergent transcription upon CTCF depletion (left), and the remaining genes with unaltered level of divergent transcription upon CTCF depletion (right). **h.** Overall transcription activity comparison between untreated cells and upon transcription inhibition. Violin plots were derived based on raw sequencing read counts representing nascent RNA molecules detected by metabolic labeling in transient-transcriptome-seq (TT-seq). **i.** Supercoiling around genes upon FLV-mediated transcription inhibition (left), with quantification of excessive negative supercoiling from two replicates (right). Based on transcription activity measured by TT-seq upon FLV treatment, high transcription genes are top 20%, low transcription genes are bottom 50%, and medium transcription genes are in between.

**Supplementary Figure 2.**
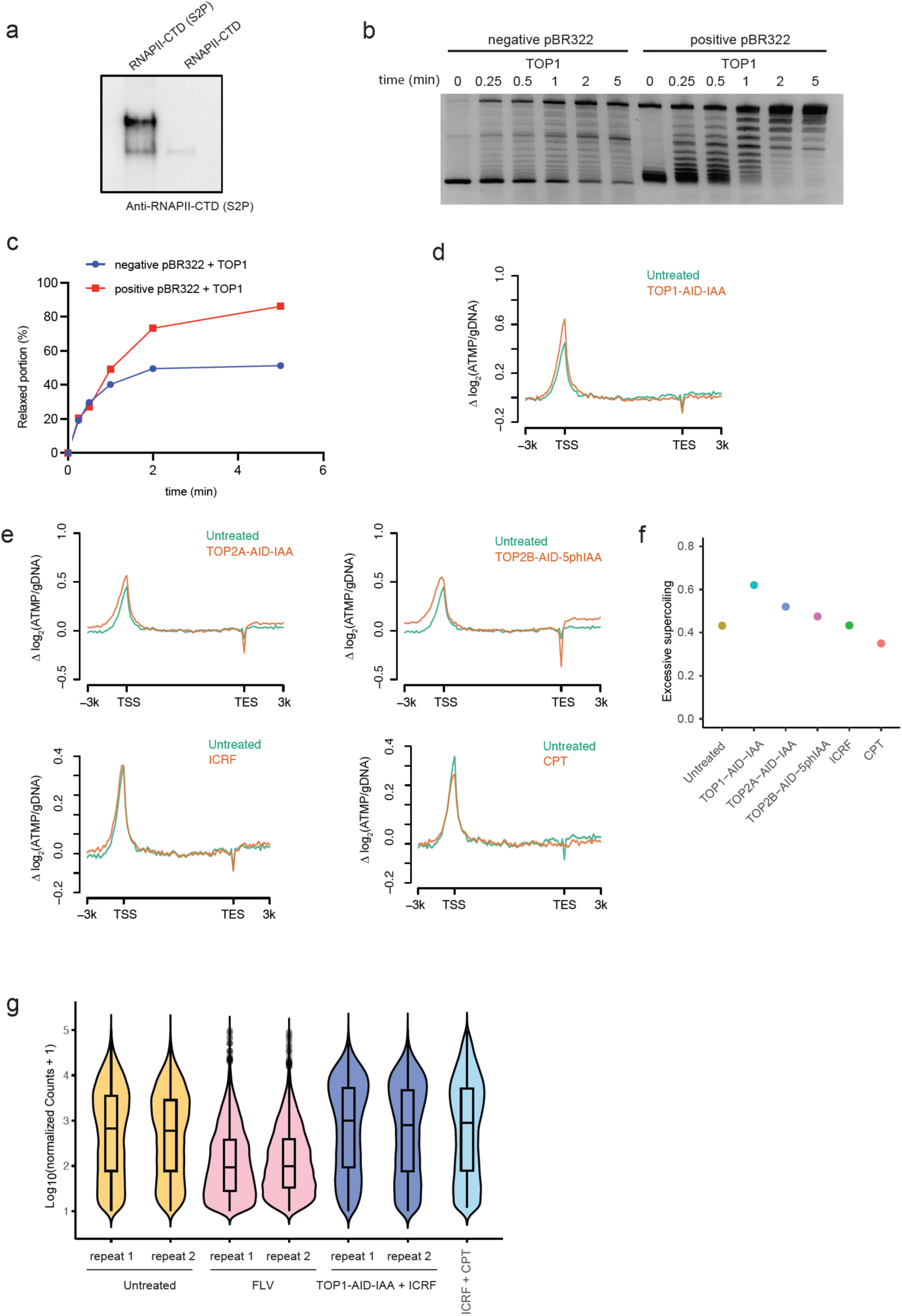
Preferential positive supercoiling relaxation during RNAP elongation. **a.** Western blot using RNAPII-CTD (S2P) antibody against 100 ng RNAPII-CTD (S2P) prepared from CycT1-cdk9-mediated in vitro kinase reaction that phosphorylates RNAPII-CTD. **b.** Negatively and positively supercoiled pBR322 were relaxed by human TOP1, with remaining supercoiling level at indicated time points quantified by agarose gel electrophoresis. **c.** Negative and positive supercoiling relaxation rate was quantified based on the gel in (b), with normalization to time point 0. **d.** Supercoiling around genes in untreated cells and upon TOP1 AID depletion. **e.** Supercoiling around genes in untreated cells and upon TOP2A or TOP2B AID depletion (top), or upon TOP2 or TOP1 drug inhibition (bottom). **f.** Quantification of excessive negative supercoiling under various conditions in (d) and (e). **g.** Overall transcription activity comparison between untreated cells, transcription inhibition conditions, and simultaneous TOP1 and TOP2 perturbation conditions. Violin plots were derived based on raw sequencing read counts representing nascent RNA molecules detected in TT-seq.

**Supplementary Figure 3.**
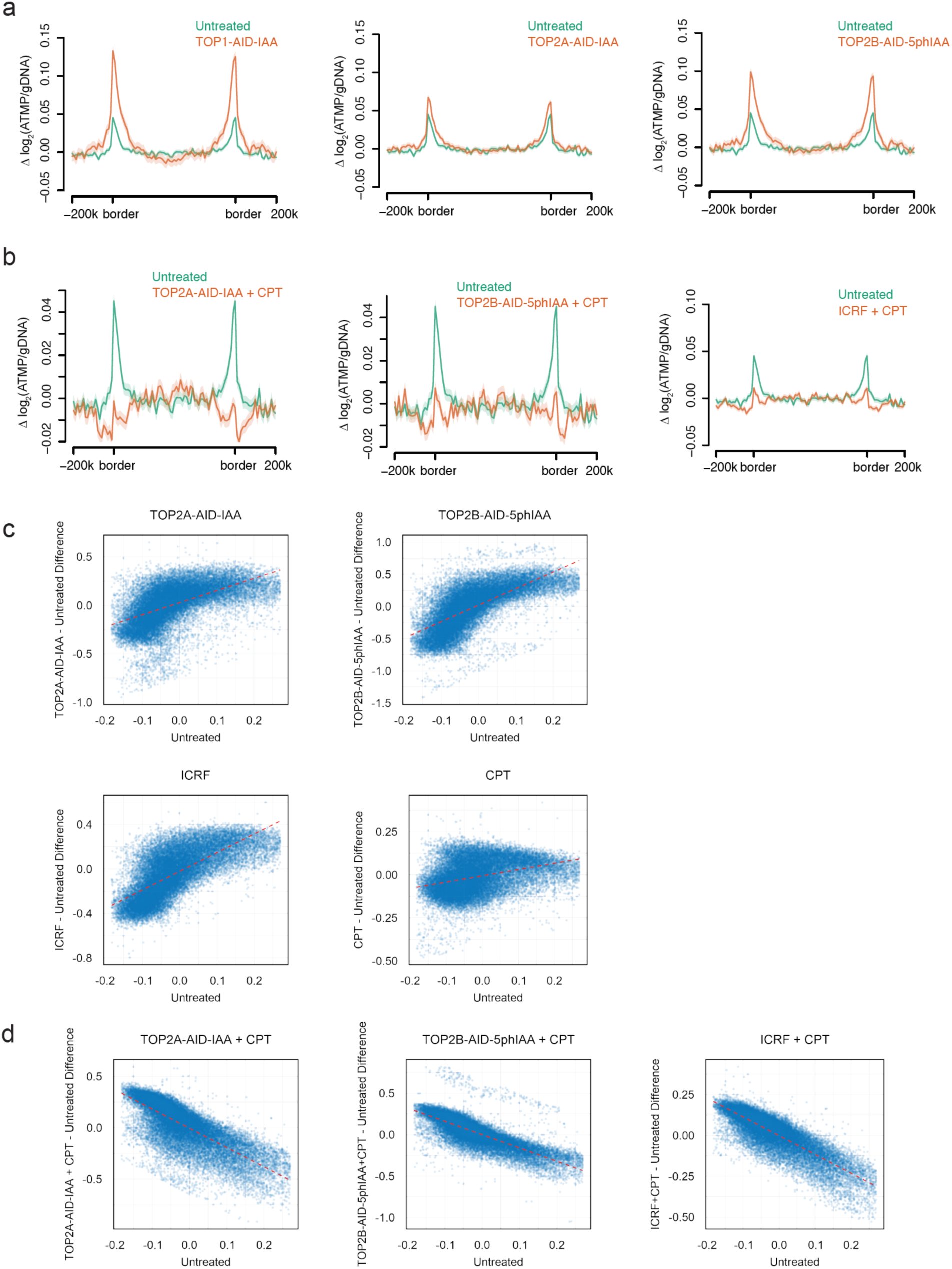
Transcription dependence of large-scale genome-wide supercoiling. **a.** Supercoiling at TAD boundaries in untreated cells and upon topoisomerase AID depletions. **b.** Supercoiling at TAD boundaries in untreated cells and under conditions of simultaneous TOP1 and TOP2 perturbation. **c.** Changes in genome-wide supercoiling under conditions of topoisomerase AID depletion or drug inhibition. Dotted lines indicate general elevation in supercoiling levels. Bin size: 100 kb. **d.** Changes in genome-wide supercoiling under conditions of simultaneous TOP1 and TOP2 perturbation. Dotted lines indicate general reduction in supercoiling levels. Bin size: 100 kb.

**Supplementary Figure 4.**
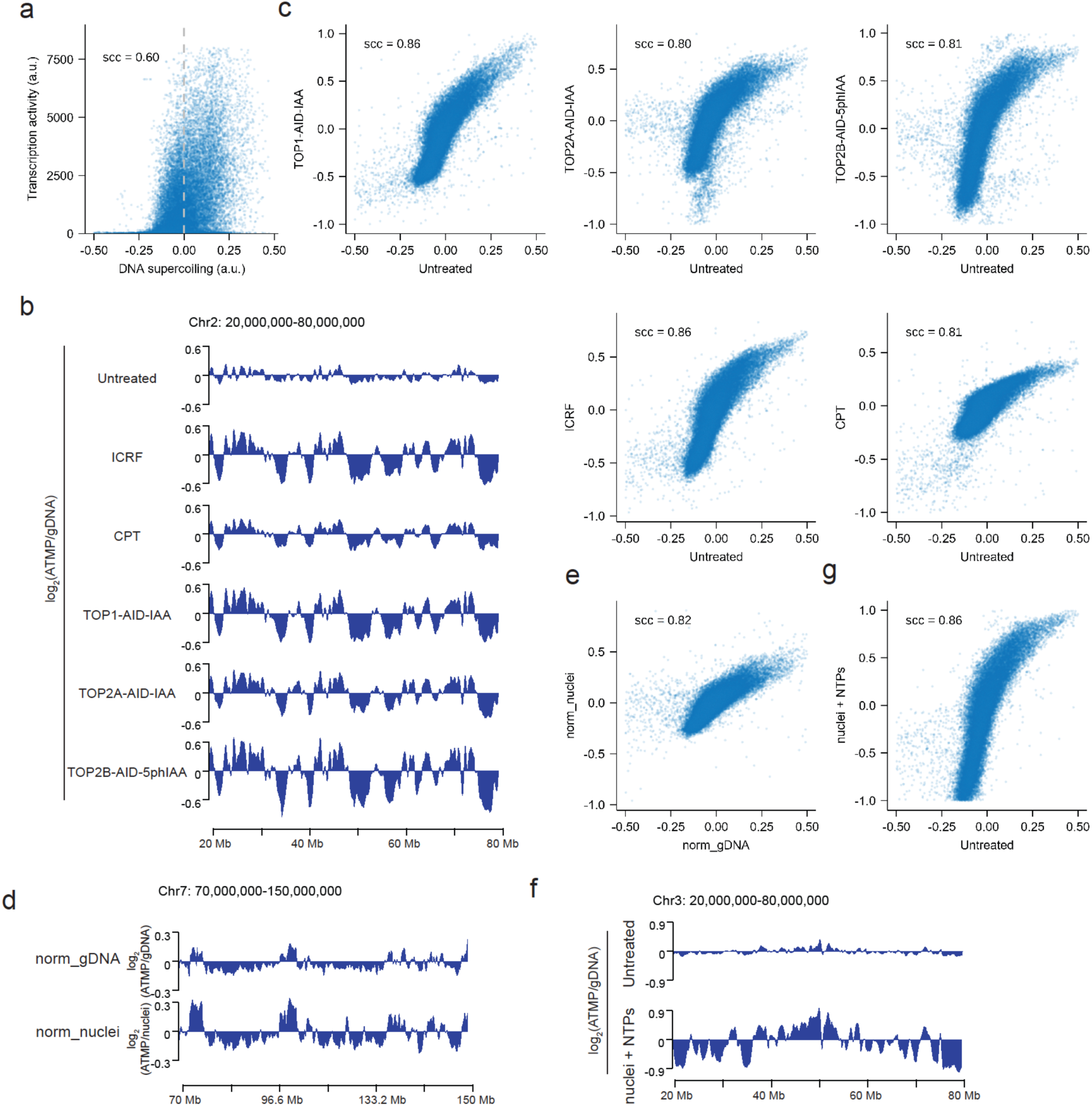
Active chromatin processes other than transcription contribute to genome-wide supercoiling. **a.** Correlation between genome-wide distributions of DNA supercoiling and transcription activity (TT-seq of nascent RNA via metabolic labeling). Bin size: 10 kb. **b.** Example regions of genome-wide supercoiling distribution under conditions of untreated and various topoisomerase perturbations. Bin size: 100 kb. **c.** Correlation of genome-wide supercoiling distributions between the untreated cells and various topoisomerase perturbation conditions. Bin size: 100 kb. **d.** Example regions of genome-wide supercoiling distribution using two different normalization methods: normalized by gDNA and normalized by extracted nuclei. Bin size: 100 kb. **e.** Correlation of genome-wide supercoiling distributions between gDNA normalization and extracted nuclei normalization. Bin size: 100 kb. **f.** Example regions of genome-wide supercoiling distribution in untreated cells and extracted nuclei with NTP addition. Bin size: 100 kb. **g.** Correlation of genome-wide supercoiling distributions between untreated cells and extracted nuclei with NTP addition. Bin size: 100 kb.

**Supplementary Figure 5.**
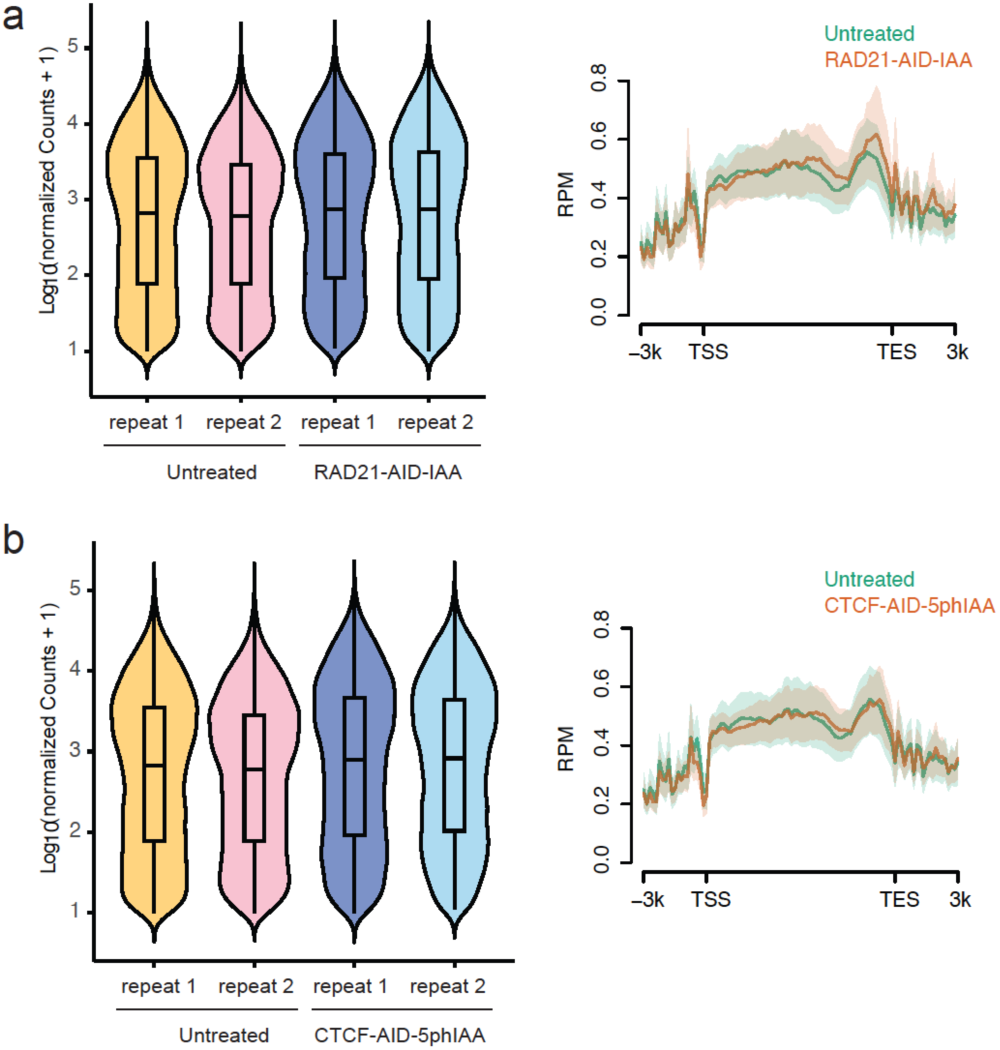
Transcription activity measurement upon cohesin or CTCF depletion. **a.** Overall transcription activity comparison between untreated cells and cohesin AID depletion condition. Violin plots were derived based on raw sequencing counts representing nascent RNA molecules detected in TT-seq (left), and nascent RNA sequencing reads per million were mapped across the gene body (right). **b.** Overall transcription activity comparison between untreated cells and CTCF AID depletion condition. Violin plots were derived based on raw sequencing counts representing nascent RNA molecules detected in TT-seq (left), and nascent RNA sequencing reads per million were mapped across the gene body (right).

**Supplementary Figure 6.**
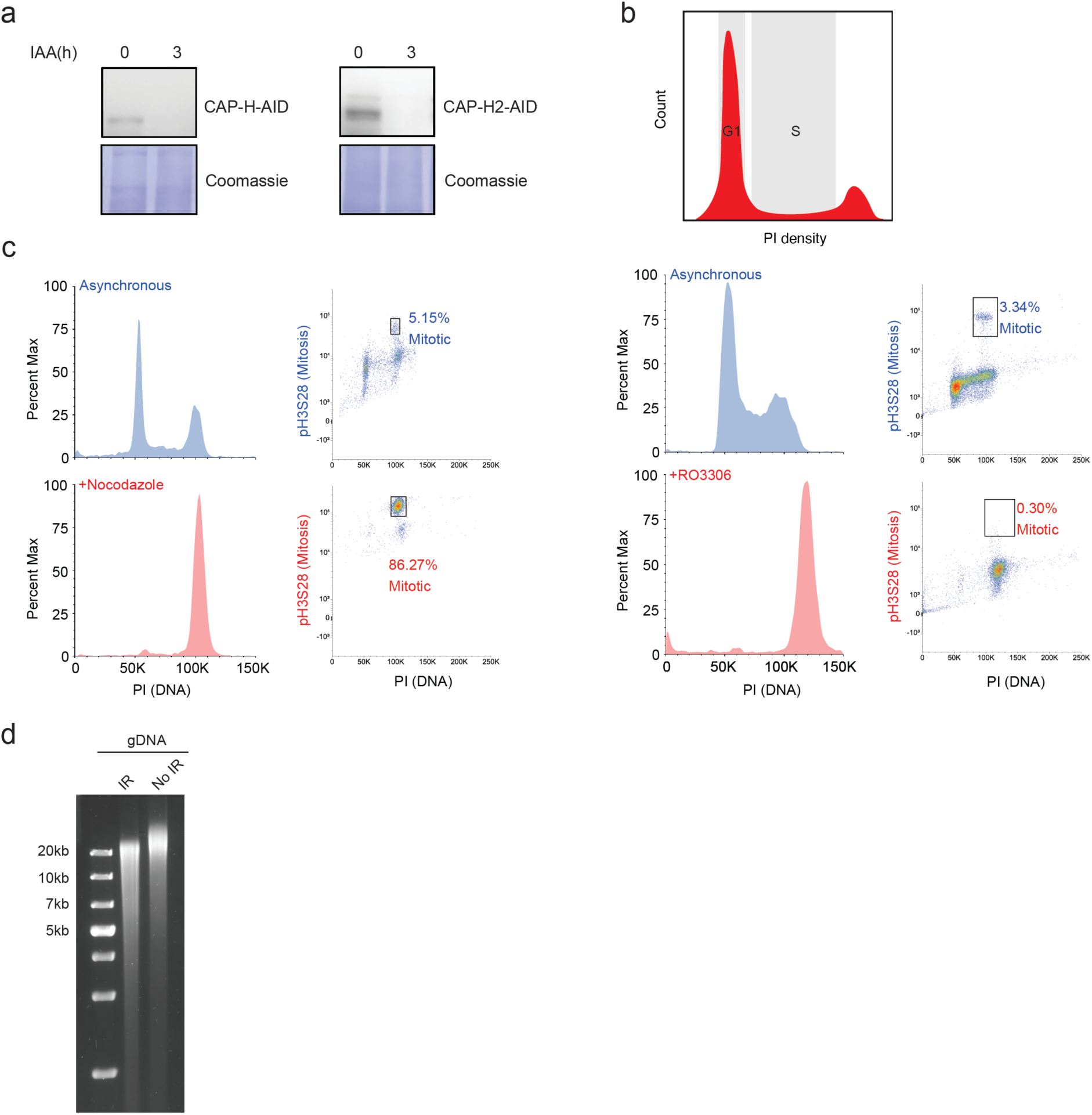
Condensin depletion, cell synchronization, and gamma radiation. **a.** Western blot showing AID depletions of condensin I subunit CAP-H and condensin II subunit CAP-H2, in HCT116 cells synchronized at prometaphase. **b.** Flow cytometry gating to sort cells at G1 phase and S phase. **c.** Flow cytometry gating to validate cell synchronization at prometaphase by nocodazole treatment, and at the G2/M checkpoint by RO-3306 treatment before drug release into early prophase. **d.** Pulsed-field gel electrophoresis (PFGE) of extracted genomic DNA from human cells with or without gamma irradiation treatment. Extracted gDNA were treated with nuclease S1 to convert single-stranded DNA breaks into double-stranded DNA breaks before PFGE analysis.

**Supplementary Figure 7.**
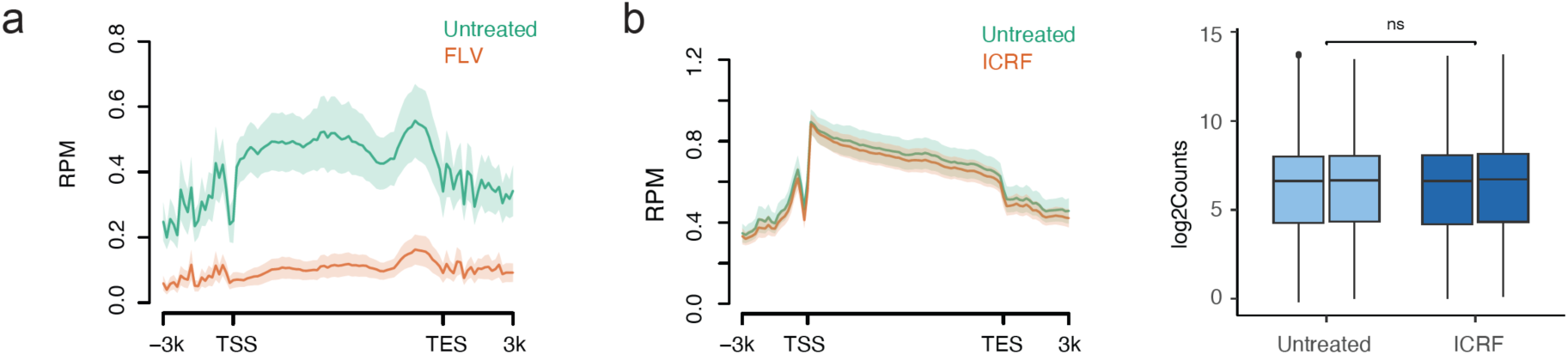
Negative supercoiling constrains human transcription activity. **a.** Transcription activity across the gene body, compared between conditions of untreated and FLV treatment. **b.** Transcription activity across the gene body, compared between conditions of untreated and ICRF treatment (left), and biological duplicates (right).

**Supplementary Figure 8.**
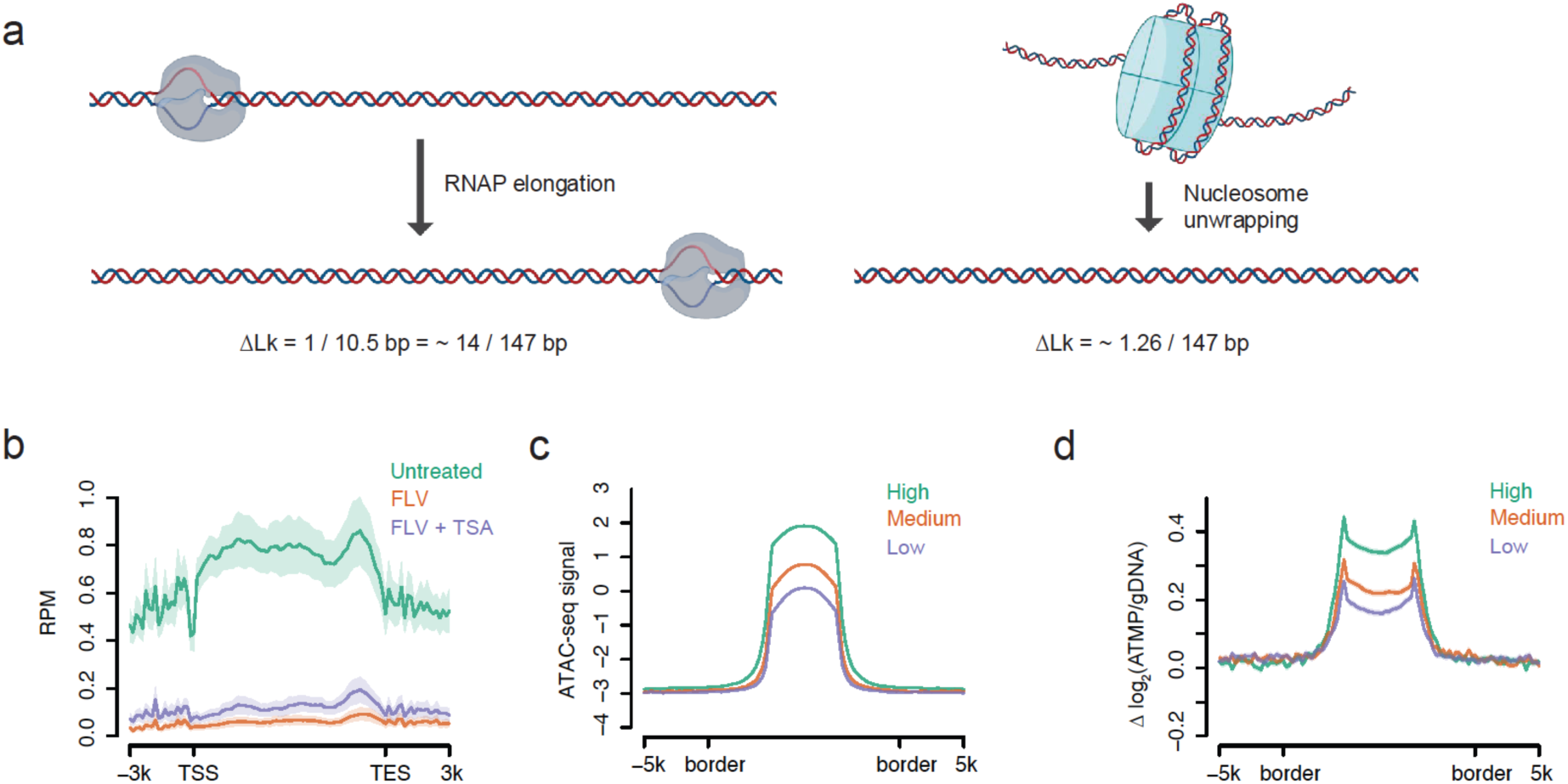
Role of nucleosomes in genome-wide supercoiling. **a.** Schematic and quantitative estimation of supercoiling generated by RNAP elongation and by nucleosome turnover. **b.** Transcription activity across the gene body under indicated conditions. **c.** ATAC-seq signal at accessible chromatin regions categorized by ATAC-seq signal. High regions are top 20%, low regions are bottom 50%, and medium regions are in between. Bin size: 200 bp. **d.** Supercoiling elevation upon TSA treatment under transcription-inhibited conditions, at accessible chromatin regions categorized by ATAC-seq signal. High regions are top 20%, low regions are bottom 50%, and medium regions are in between. Bin size: 200 bp.

## Methods

### Cell culture

HCT116 cell line was cultured in McCoy’s 5A medium (ThermoFisher, 16600108) supplemented with 10% fetal bovine serum (FBS) at 37 °C under 5% CO_2_ in the incubator. GM12878 cell line was cultured in RPMI 1640 medium (Corning, 15-040-CM) supplemented with 2 mM l-glutamine (Corning, 25-005-CI) and 15% FBS at 37 °C under 5% CO_2_ in the incubator. eHAP, a fully haploid human cell line derived and modified from chronic myeloid leukemia, was cultured in Iscove’s Modified Dulbecco’s Medium (IMDM) supplemented with 10% FBS at 37 °C under 5% CO_2_ in the incubator. *Drosophila* S2 cell line was cultured in Schneider’s *Drosophila* medium supplemented with 10% heat-inactivated FBS at 27 °C in the incubator. Heat-inactivated FBS was generated by heating FBS in 56 °C water bath for 60 min. All experiments used HCT116 cells unless specified otherwise.

### Yeast strains, culture methods, and treatment

The *Saccharomyces cerevisiae* strain CH1585 were grown overnight to achieve OD between 0.2∼0.3 in YPD medium at 26 °C. To synchronize the cells, alpha-Factor (T6901; Sigma) was added to a final concentration of 5 µg/ml for 1 h. A second dose of alpha-Factor (5 µg/ml) was then added, followed by another 1.5 h incubation. Following synchronization, cells were harvested and washed with PBS and resuspended in 1 ml of PBS. Cells were then incubated with 100 µM of ATMP with 5 µg/ml of alpha-factor for 5 min at 26 °C with shaking, followed by UV-A irradiation at 360 nm for 1 min. Immediately after irradiation, NaN3 was added to a final concentration of 10 mM. Cells were subsequently resuspended in 600 µl of sorbitol buffer, and 10 µl Lyticase (L2524; Sigma) was added. Samples were incubated at 30 °C for 30 min. Cells were centrifuged at 300 × g for 10 min and resuspended in 500 µl TE buffer. Subsequently, 50 µl of 10% SDS, 5 µl RNase A and 10 µl of Proteinase K were added, and samples were incubated at 56 (C for at least 4 h. Samples were cooled on ice and 550 µl Phenol-Chloroform-isoamyl alcohol (25:24:1: sigma) was added. The samples were vortexed for 20 s and transferred to phase-lock tube, followed by centrifugation at 16, 000 × *g* for 10 min. The aqueous phase was further extracted with 300 µl of Chloroform by gently inverting 10 times, followed by centrifugation at 16, 000 × *g* for 3 min. The aqueous phase was transferred to a new microcentrifuge tube without disturbing the interphase. Nucleic acids were precipitated with 1 ml ice-cold ethanol, washed with ice-cold 70% ethanol two times, and dissolved in TE buffer. For the gDNA control, cells synchronized by the same two-step alpha-Factor treatment were subjected to Lyticase digestion without ATMP treatment. The extracted gDNA was sonicated to 600 bp and subsequently incubate with ATMP and exposed to the same UV-A treatment to achieve an equivalent *in vitro* adduct density. Subsequent steps were performed as described for the human samples.

### Cell culture perturbation conditions

Transcription initiation was inhibited by adding triptolide (TPL) to the cell culture medium to a final concentration of 1 µM for 1-h incubation. Transcription elongation was inhibited by adding flavopiridol (FLV) to the cell culture medium to a final concentration of 1 µM for 45-min incubation. HDAC inhibition was done by adding trichostatin A (TSA) to the cell culture medium to a final concentration of 500 nM for 24-h incubation. BAF complex was inhibited by adding BRM014 (BRM) to the cell culture medium to a final concentration of 1 µM for 6-h incubation. Target protein depletions in different cell lines via the AID system were performed following conditions in our previous work [1]. Briefly, TOP1 and TOP2 were inhibited by a 10-min incubation with 10 µM camptothecin (CPT) (Sigma, 208925) and 10 µM ICRF-193 (ICRF) in the cell culture medium, respectively. For AID depletion of TOP1 and TOP2A, the corresponding cell lines were incubated with 500 µM indole-3-acetic acid (IAA) (Sigma, I3750) in the medium at 37 °C for 2 h. For AID depletion of RAD21, the corresponding cell line was incubated with 500 µM IAA in the medium at 37 °C for 6 h. For AID2 depletion of CTCF and TOP2B, the corresponding cell lines were incubated with 1 µM 5-ph-IAA (MedChemexpress, HY-134653) in the medium at 37 °C for 4 h.

### Cell sorting, synchronization, and perturbation

We used different strategies for synchronizing and enriching cells at different cell cycle stages. For G1 phase and S phase cells, HCT116 cells were fixed by cold 70% ethanol, treated with RNase A, and stained with PI. Cells in G1 and S cell-cycle stage were gated based on DNA content. For cells at prometaphase or prophase stages, cells were blocked in S phase by the addition of 2.5 mM thymidine (Sigma, T9250) for 22-24 hours. After washing with PBS (ThermoFisher, 10010049) three times, the cells were released into fresh media for 4 hours. Cells were then enriched to prometaphase by the addition of 400 ng/mL nocodazole (Sigma, M1404) for 14 hours or to the early prophase by the addition of 9 µM RO-3306 (Sigma, SML0569) for 12 hours followed by drug release. Cell lines expressing CAP-H-mAID-mCherry or CAP-H2-mAID-mCherry (Ref) were treated with 0.5 mM IAA (Sigma, I3750) during the final 3 hours of nocodazole treatment to deplete AID tagged proteins.

### Flow cytometry analysis

Cells were harvested with trypsin (ThermoFisher, 25200056) and washed with ice-cold PBS before fixation with –20 °C 75% EtOH. Cells were washed twice with ice-cold PBS before a final wash using ice-cold PBSB (PBS + 1% BSA). For immunostaining, cells were resuspended in ice-cold PBSBT (PBS + 1% BSA + 0.1% Triton-X) and incubated with pH3S28 antibody (ab237418, dilution 1:500) for 1 hour at room temperature. After two washes with ice-cold PBSBT, cells were stained with 0.1 mg/mL propidium iodide (ThermoFisher, P3566) and RNA was digested by 0.1 mg/mL RNase A (ThermoFisher, EN0531) for 30 minutes to 1 hour at room temperature. Unstained and single stained samples served as controls. FCS, SSC, PI fluorescence, and Alexa 647 fluorescence were collected on a BD LSRFortessa Cell Analyzer. Data was analyzed using Floreada.io. Single cells were gated using FCS and SSC values. Mitotic cells were gated based on pH3S28 staining, and the percentage of mitotic cells was calculated.

### In vitro kinase reaction to prepare RNAPII-CTD (S2P)

Serine 2 phosphorylation of human RNAPII-CTD was generated by incubate human RNAPII-CTD protein (Abcam, ab81834) with CycT1-cdk9 (a gift by Mardo Koivomagi lab) in a kinase reaction containing 50 mM Tris-HCl pH 7.5, 5 mM DTT, 5 mM MnCl2, 4 mM MgCl_2_, and 1 mM ATP. After the kinase reaction, we purified RNAPII-CTD (S2P) by following instructions of Pierce™ Glutathione Magnetic Agarose Beads (Thermo Scientific, 78601) with minor modifications, since the original human RNAPII-CTD protein contain a GST tag. The purified RNAPII-CTD (S2P) was stored at –80°C with 20% glycerol. The S2P phosphorylation was verified by Western blot using RNAPII-CTD S2P antibody (Abcam, ab5095).

### In vitro topoisomerase relaxation assay and DNA supercoiling gel electrophoresis

5 ng of recombinant human TOP1 (Abcam, ab82099) was incubated with 10 ng RNAPII-CTD or RNAPII-CTD (S2P) for 10 min on ice in the TOP1 buffer (10 mM Tris-HCl pH 7.5, 10 mM MgCl_2_, 150 mM NaCl, 0.1 mM EDTA, 15 ug/ml BSA). After the addition of 500 ng plasmid DNA (positively supercoiled PBR322 (inspiralis, POS5001) or negatively supercoiled PBR322 (inspiralis, S5001)), reactions were incubated at 37°C for different time durations and terminated by the addition of 1% TE-SDS and 200 µg/ml proteinase K. Purified DNA products were analyzed on a 1% (w/v) agarose gel. To visually separate different topoisomer species with different supercoiling level, gels were run in TAE buffer at 40 V for 16 hours. Gels were then stained with GelRed in 1X TAE buffer for 1 hour and imaged with Bio-Rad ChemiDoc MP Imaging System. The quantification of DNA supercoiling level based on topoisomer bands on the gel was the same as before^1^, and topoisomerase relaxation curve was subsequently determined for each incubation condition based on supercoiling level at different time points.

### Nuclei extraction

Nuclei from human cells were prepared following the liquid Hi-C protocol [2], with chromatin structures remain intact in the extracted nuclei. Started from 50 million HCT116 cells, the extracted nuclei were dissolved in an adequate total volume to obtain 2 million nuclei per 0.2 ml nuclei storage buffer (10 mM PIPES pH 7.4, 10 mM KCl, 2 mM MgCl_2_, 50% glycerol, 8.5% sucrose, 1 mM DTT (added before use), 1:100 protease inhibitor (added before use)). For ATMP-seq, nuclei were washed twice with nuclear isolation buffer (10 mM PIPES pH 7.4, 10 mM KCl, 2 mM MgCl_2_, 1 mM DTT (add before use), 1:100 protease inhibitor (add before use), pH adjusted to 7.4 using 1 M KOH) to remove glycerol, and resuspend with nuclear isolation buffer for ATMP incubation. For nuclei with the addition of NTPs, nuclei were resuspended with nuclear transcription run-on buffer (10mM Tris-HCl pH 8, 5mM MgCl_2_, 300mM KCl, 1mM DTT, 500 uM each NTPs) and incubate at 30°C for 10 min, followed by wash with nuclei isolation buffer, before proceeding with the same protocol for ATMP incubation for extracted nuclei. For TOP1 treatment, the nuclei were resuspended with TOP1 buffer (50 mM Tris-HCl pH 7.5, 50 mM KCl, 10 mM MgCl_2_, 0.1 mM EDTA, 0.5 mM DTT, 30 μg/ml BSA) with 100 unit of Topoisomerase I (ThermoFisher, 38042024) and incubate at 37°C for 1 hour, followed by wash with nuclei isolation buffer, before proceeding with the same protocol for ATMP incubation for extracted nuclei.

### ATMP-seq

ATMP-seq was carried out according to a previously established protocol [1], with minor modifications. In brief, cultured cells were exposed to 100 μM Azide-TEG-TMP (ATMP; Berry and Associates, PS 5030) in growth medium for 10 min at 37 °C inside the incubator. This was immediately followed by UV-A irradiation at 360 nm for 10 min, delivering a total dose of 6 kJ/m² and producing an ATMP adduct density of approximately one monoadduct per 10 kb. Following irradiation, cells were fixed in ice-cold 75% ethanol for 30 min, rinsed three times with PBS, and treated with 800 μg/ml RNase A for 1 h. Genomic DNA was isolated using the DNeasy Blood & Tissue Kit (Qiagen, 69504) with RNase A digestion, and then fragmented to ∼600 bp using a Covaris M220 ultrasonicator. For each 1 μg of DNA, 1 μl of 10 mM DBCO-desthiobiotin was added, and the mixture incubated at 37 °C for 2 h with intermittent shaking at 950 rpm. DNA was then cleaned up using the MinElute PCR Purification Kit (Qiagen, 28004). Biotinylated fragments were captured with Dynabeads MyOne Streptavidin T1 magnetic beads (ThermoFisher, 65601), and DNA was released from the beads in an elution buffer containing 50 mM Tris (pH 8), 200 mM NaCl, and 10 mM biotin, incubated for 1 h at room temperature with rotation. The recovered DNA was quantified using the Qubit dsDNA HS assay, and ATMP density was determined from the ratio of input DNA to recovered DNA. For sequencing, eluted DNA was further purified and sheared to ∼150 bp, followed by library preparation using the NEBNext Ultra II DNA Library Prep Kit for Illumina. Libraries were sequenced to a depth of ∼40 million reads (either 1 × 50 bp on a NextSeq 2000 or 1 × 75 bp on an AmpSeq platform). For the genomic DNA normalization control, extracted gDNA was first fragmented to ∼600 bp, then incubated with ATMP and exposed to the same UV-A treatment to achieve an equivalent in vitro adduct density. Subsequent steps mirrored those used for the experimental samples.

### TT-seq

TT-seq was conducted based on a previously described protocol [3] with slight modifications. Briefly, cells were cultured for 24 h and subsequently labeled with 500 μM 4-thiouridine (4sU; ThermoFisher, J60679-MD) for 20 min in growth medium. Immediately after labeling, RNAzol (Sigma, R4533) was added to lyse the cells. Total RNA was isolated according to the manufacturer’s instructions, and 400 ng of 4sU-labeled *Drosophila* spike-in RNA (prepared by incubating with 500 μM 4sU for 2 h) was added per 100 μg of extracted RNA. The RNA was fragmented to an average length of ∼1 kb using a Covaris M220 sonicator. Thiol-containing RNA was biotinylated by incubation with Biotium Biotin-XX MTSEA (ThermoFisher, 50-196-4817) at room temperature for 30 min in the dark. Following biotinylation, total RNA was cleaned with phenol–chloroform–isoamyl alcohol extraction, and the labeled RNA was captured using μMACS streptavidin beads (Miltenyi Biotec, 130-074-101). Bound RNA was eluted with 100 mM DTT. The recovered nascent RNA was treated with DNase I, purified using the RNA Clean & Concentrator kit (Zymo Research, R1014), and used for library preparation with the NEBNext rRNA Depletion Kit v2 (NEB, E7405) and NEBNext Ultra II RNA Library Prep Kit (NEB, E7490). Libraries were sequenced on an Illumina platform for about ∼40 million reads (either 1 × 75 bp or 1 × 75 bp on an AmpSeq platform).

### Validation of target protein depletion by Western blot

Cells were washed with ice-cold PBS and lysed using 1x sample buffer (62.5 mM Tris pH 6.8, 10% glycerol, and 2% SDS). Cell lysates were boiled at 95 °C for 30 minutes and protein concentrations in each sample were determined using the DC Protein Assay kit (Bio-Rad). Lysates were then diluted to 2 µg/µL of protein using 1x sample buffer, before the addition of 50 mM DTT and 0.005% bromophenol blue. Before western blotting, the final lysates were boiled at 95 °C for 5 minutes. Equal amounts of protein from each sample were loaded in a 6% hand-casted SDS-PAGE gel (Bio-Rad, 1610158) for electrophoresis, followed by transfer to a LF-PVDF membrane (Bio-Rad, 1620261) using the Mini Trans-Blot Cell system (Bio-Rad). LF-PVDF membranes were then blocked by 5% nonfat milk for 1 h and incubated overnight at 4 °C with primary antibody against CAP-H (Proteintech, 11515-1-AP, 1:1000 dilution) or CAP-H2 (Proteintech, 26172-1-AP, 1:1000 dilution). After TBST washes, membranes were incubated with anti-rabbit secondary antibody (Abcam, ab6721, 1:10,000 dilution) for 1 hour at room temperature and followed with additional TBST washes. The target proteins were visualized using the SuperSignal West Pico PLUS chemiluminescent substrate (ThermoFisher), and the images were captured using the ImageQuant 800 biomolecular imager (Cytiva). After imaging, membranes were stripped using stripping buffer (0.2 M glycine, pH 2.2, 0.1% SDS, 1% Tween-20) for 5 mins at room temperature. Membranes were then rinsed with TBST and stained with Coomassie Brilliant Blue R-250 to visualize total protein as loading control.

### Gamma irradiation

Cells were aliquoted in a 6-cm culture dish and placed on ice while irradiated by the Mark-1 γ-irradiator (JL Sherpherd and Associates) for a total dosage of 500 Gy. Cells for unirradiated control were taken into the same irradiation room.

### Pulsed-field gel electrophoresis (PFGE)

PFGE was performed using the BioRad CHEF-DR II system. 0.2-2.2 Mb S. cerevisiae ladder (BioRad, 1703605) was used as the high molecular weight reference. Ladders and samples were loaded on the 1% agarose gel in 0.5X TBE buffer. Electrophoresis was carried out with 60 to 120 seconds ramped pulse times at 6 V/cm for 24 hours. Temperature was controlled at 14°C throughout the electrophoresis.

### Genome-wide detection of TOP2A and TOP2B cleavage complexes

Seven to ten million cells were washed once with ice-cold 1× PBS and immediately transferred to an ice-cold water bath. Genomic DNA was purified using DNAzol™ Reagent according to the manufacturer’s instructions. Following ethanol washing, the DNA pellet was briefly air-dried and dissolved in 100 μl of 1× TE (10 mM Tris-HCl pH 8.0, 1 mM EDTA) containing 1% SDS, and incubated at room temperature for 10 min. The sample was then supplemented with 100 μl of 1× TE containing 100 ng/μl RNase A and incubated at room temperature for 10–20 min, followed by addition of DTT to a final concentration of 10 mM and incubation at room temperature for another 10 min. The mixture was brought to 600 μl with nuclease-free water, and DNA was precipitated by adding an equal volume of 2-propanol and 1/10 volume of 3 M sodium acetate, followed by centrifugation at 4,000 × g for 5 min to pellet the DNA. The pellet was washed with 70% ethanol and dissolved in 1× TE containing 0.5% SDS. DNA concentration was measured using the Qubit dsDNA Broad Range Assay Kit.

Sixty micrograms of genomic DNA was sonicated to approximately 500 bp using a Covaris M220 (130 μl AFA microtube) according to the manufacturer’s settings. The sonicated DNA was diluted to 200 μl with ingredients to meet the concentration of 1× IP buffer (1× TE, 1× PBS, 1% Triton X-100). A 300 μl IP mixture was then prepared with 1× IP buffer containing 1–2 μg of antibody (anti-TOP2A or anti-TOP2B) and spike-in chromatin with corresponding spike-in antibody, which could be ordered from Active Motif; a pilot assay should be performed to determine the optimal amount needed. Twenty microliters of the mixture was reserved as input control. The IP mixture was rotated overnight at 4°C.

The following day, 10–20 μl of Dynabeads Protein G/A was added to the IP mixture and incubated with rotation for 3 h at 4°C. Beads were washed sequentially with 400 μl of 1× IP buffer (2 washes, 7 min each with rotation), high-salt wash buffer (50 mM Tris-HCl pH 8.0, 500 mM NaCl, 1 mM EDTA, 1% Triton X-100, 0.1% SDS; 2 washes, 7 min each), LiCl wash buffer (10 mM Tris-HCl pH 8.0, 250 mM LiCl, 1 mM EDTA, 1% NP-40, 1% sodium deoxycholate; 2 washes, 7 min each), and 1× TE (2 washes, 5 min each), transferring beads to a new tube at the final wash step. DNA was eluted from the beads in 30 μl 1× TE containing 100 ng/ml proteinase K, incubated at 55°C for 2 h, followed by denaturation at 95°C for 10 min.

The eluate was then treated with 0.5 μl of TDP2 in 1× TDP2 reaction buffer at final volume of 55 μl for 1 h to remove residual covalently bound topoisomerase peptide from the DNA 5′ ends. The TDP2-treated DNA was sonicated to approximately 250 bp using a Covaris M220 (55 μl AFA microtube) and purified using 2.5 volume of DNA purification beads (e.g., AMPure XP or equivalent). Sequencing libraries were constructed using the Vazyme ssDNA Library Prep Kit or equivalent kit from IDT according to the manufacturer’s instructions.

### Sequencing data analysis

For ATMP-seq, raw sequencing reads were quantity-trimmed using Trimmomatic [4] and aligned to the hg38 reference genome with Bowtie 2 [5]. The resulting alignments were sorted and indexed using SAMtools [6] and deduplicate reads were removed with Picard. Normalized coverage tracks in bigWig files were generated with deepTools [7], employing the bamCoverage and bigwigCompare functions. Supercoiling profiles for gene bodies and TAD regions were generated using ngsplot [8], normalized either to relaxed supercoiling level in gene bodies or to non-TAD boundary regions. Correlation analyses were performed using 100 kb bins bedgraph files.

For TT-seq, adapter and quality trimming was performed with Trimmomatic, followed by alignment to hg38 and dm6 (Drosophila) reference genome by STAR [9] with the parameters –-outFilterMismatchNoverReadLmax 0.02 –-outFilterMultimap ScoreRange 0 –-alignEndsType EndToEnd. Alignment results to the dm6 genome (spike-in control) were used for cross-sample normalization [3]. The resulting sorted bam files were used to produce gene body profile via ngsplot.

